# LIMBER: A Hybrid RNA Foundation Model for Predicting and Understanding RNA Structure and Function

**DOI:** 10.64898/2026.10.28.762670

**Authors:** Ziyao Wang, Yunrui Qiu, Yexiao He, Isabel Thompson, Hongli Wang, Adam Porter, Ang Li

## Abstract

RNA foundation models offer a promising route to decoding the relationships between RNA sequence, structure and function, yet how model architecture shapes the acquisition and organization of biological information remains poorly understood. Here we introduce Layer-Interleaved Mamba-Bidirectional Encoder for RNA (LIMBER), a hybrid foundation model integrating bidirectional self-attention with selective state-space modeling, pre-trained on 40.5 million non-coding RNA sequences. With task-specific fine-tuning, LIMBER achieves leading performance on secondary-structure prediction and non-coding RNA classification benchmarks, while demonstrating strong transferability to translation-related tasks and RNA–small-molecule interaction prediction. Beyond model development, we establish a unified, task-agnostic framework to systematically evaluate and interpret the biological knowledge intrinsically acquired by RNA foundation models. This framework validates LIMBER’s strong capabilities in recovering RNA structures and predicting mutational effects, while uncovering a hierarchical, task-dependent organization of biological information across model layers. In addition, we demonstrate distinct scaling behaviors across structural, evolutionary, and functional capabilities by training and evaluating a broad panel of model variants spanning model capacity, training duration, and dataset size. Taken together, these findings establish LIMBER as a versatile RNA foundation model and provide insight into how architectural design, internal representations and scaling shape the learning of RNA sequence-structure-function relationships.

## 1 Introduction

Ribonucleic acid (RNA) is a class of polymeric biomolecules that function as both carriers of genetic information and mediators of biological processes. Only a small fraction of cellular RNAs serve as templates for protein synthesis (mRNA), however, heterogeneous families of non-coding RNAs (ncRNA) [1], including ribosomal RNA (rRNA), transfer RNAs (tRNA), microRNAs and ribozymes, are transcribed from non-protein-coding genomic regions. These ncRNAs play central roles in molecular recognition, gene regulation, and biochemical catalysis[2, 3], and are increasingly recognized as disease biomarkers and therapeutic targets[4, 5]. The diverse biochemical activities of these functional entities arise from their distinct structures, which are encoded by nucleotide sequences. Elucidating the hierarchical relationships between sequence, structure, and function for ncRNAs is essential for advancing our understanding of RNA biology and facilitating the development of RNA-targeted therapeutics[6, 7].

Non-coding RNA sequences are the products of an evolutionary selection process that retains signatures of nucleotide syntax, structural propensities, and functional constraints[8]. However, extracting this information directly from raw sequences remains challenging in the absence of curated annotations. Sequencing technologies have generated vast collections of RNA sequence data, but experimentally resolved high-resolution structures and functional labels are scarce in comparison, creating a gap between sequence availability and biological data suitable for supervised learning strategies[9]. Large language models have transformed natural language processing by learning contextual representations and dependencies from massive corpora of unlabeled text[10, 11]. Extending this paradigm to biomolecular sequences offers a promising self-supervised approach for distilling coevolutionary dependencies and functional signals directly from unannotated sequence data.

Protein language models, such as the ESM family[12] and OmegaPLM[13], have demonstrated the capability to learn representations of evolutionary couplings from sequences. These representations can replace explicit multiple sequence alignments (MSAs, inputs for leading structural prediction models such as AlphaFold2/3[14, 15]) and have enabled accurate structure prediction and function inference[16]. Accordingly, a growing number of RNA language models (RLMs) have recently emerged that similarly leverage the coevolutionary signals in RNA sequences. Most existing models focus on ncRNAs and adopt bidirectional encoder representations from transformers (BERT)-based architectures. Early models, such as RNABERT[17] and RNA-FM[18], have been followed by larger-scale models, including RiNALMo[19] and the 1.6-billion-parameter AIDO.RNA[20], alongside alternative architectures based on GPT, Llama, and Mamba[21–24]. Additional strategies have incorporated RNA-specific knowledge through structure- or motif-aware architectures and masking schemes, as in RNAErnie[25], ERNIE-RNA[26], and structRFM[27]; and MSA information, as in RNA-MSM[28]. Meanwhile, the scope of pre-training has expanded from predominantly ncRNAs to broader sequence collections containing both coding and non-coding transcripts, as in Uni-RNA[29] and HydraRNA[24], as well as specialized datasets targeting plant RNAs (e.g., PlantRNA-FM[30]), untranslated regions (e.g., UTR-LM[31]), and RNA-splicing signals (e.g., SpliceBERT[32]).

Despite rapid progress in RLMs, several fundamental questions remain insufficiently explored, particularly those concerning what biological information these models acquire during pre-training and how it is encoded within their learned representations. Addressing these questions is essential for evaluating the intrinsic capabilities of RLMs and interpreting biological signals encoded by their outputs. RLM performance is typically assessed after supervised fine-tuning for specific downstream prediction tasks. However, downstream performance may not faithfully reflect the internal qualities of the pre-trained model, as the results are inherently influenced by task-specific optimization on limited, labeled data[33–35]. Interpretability presents an array of challenges as many analysis approaches, such as sparse autoencoders[36, 37], require extensive training of auxiliary models and access to additional labeled data, which remain relatively limited for RNA. These limitations highlight the need for a unified, unsupervised framework that directly probes the capabilities of pre-trained RLMs across complementary dimensions of RNA biology including structure, function, and evolution, without the need for additional fine-tuning. Such a framework would provide a principled foundation for further improving model architectures, scaling strategies, and training methodologies.

Here, we introduce LIMBER (Layer-Interleaved Mamba–Bidirectional Encoder for RNA), a fully-open hybrid ncRNA foundation model that interleaves bidirectional self-attention[10] with selective state-space modeling[11, 38] (Fig. 1, left). This architecture models direct local and long-range nucleotide interactions with sequential context propagation and bidirectional attention, providing complementary inductive biases for representing the ordered and non-local dependencies underlying ncRNA sequences. Alongside LIMBER, we establish a zero-shot (label-free), task-agnostic framework for evaluating and interpreting intrinsic model capabilities across three dimensions. We use this framework to systematically dissect the effects of dataset size, model scale, and training budget across BERT, Mamba, and hybrid model families (Fig. 1, upper right). These analyses suggest RLMs organize structural and functional information non-uniformly across model layers and exhibit different scaling behaviors in their the capacity to predict distinct biological characteristics. Across these evaluations, LIMBER consistently demonstrates strong and balanced representations across different biological dimensions compared with other RLMs.

**Fig. 1.**
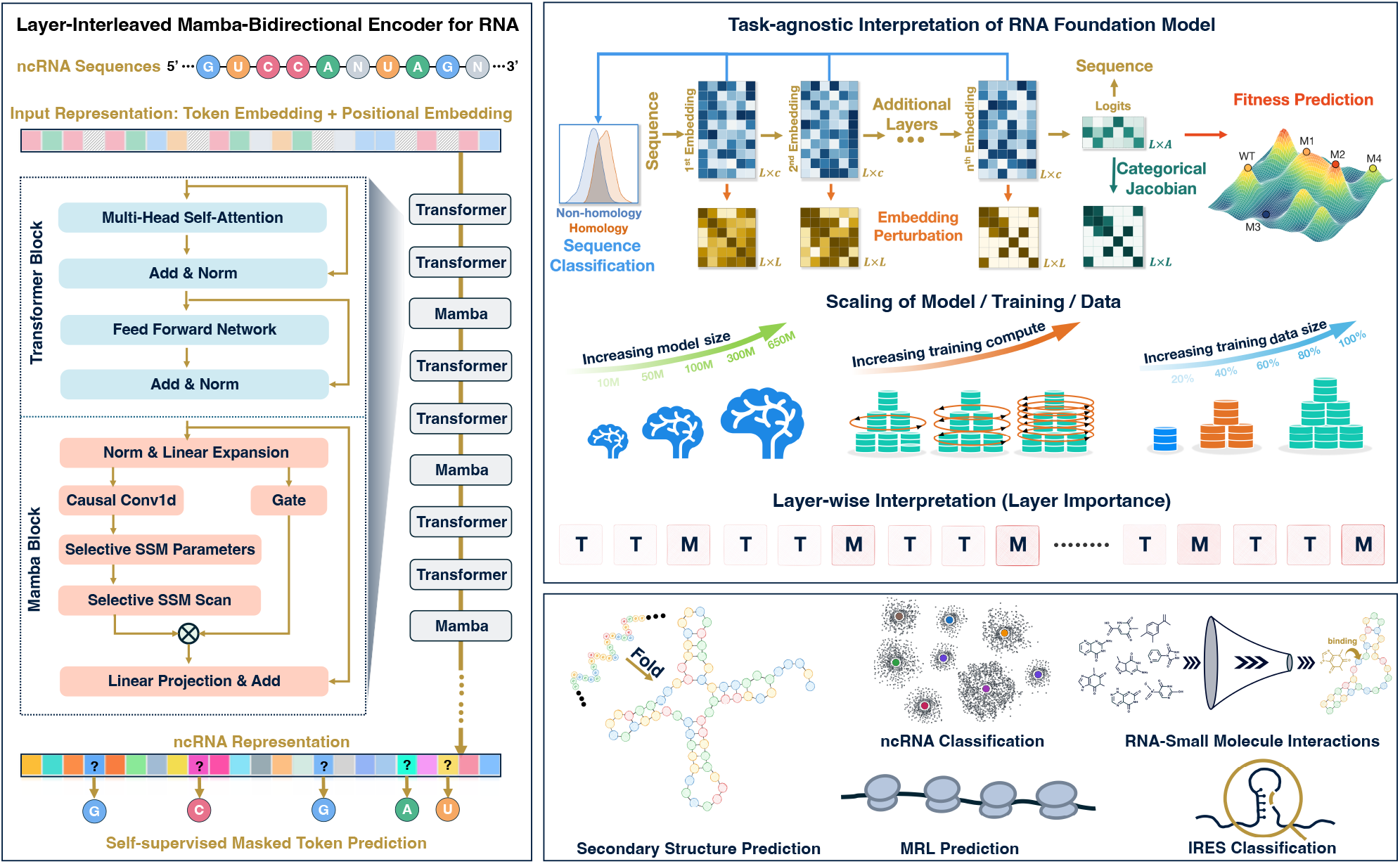
Overview of the LIMBER architecture, unified task-agnostic evaluation and interpretation framework, and downstream applications. (Left) LIMBER adopts a layer-interleaved architecture combining Mamba layers with bidirectional Transformer layers. Non-coding RNA sequences are used as inputs, and the model is pre-trained using a masked language modeling objective. (Upper right) Schematic illustration of four unsupervised strategies for evaluating and interpreting RNA language models. These methods are applied to assess RLM performance and to investigate scaling behavior with respect to model size, training compute, dataset size, and probe layer-wise importance. Secondary structure information is extracted directly and without fine-tuning from model predicted logits with the categorical Jacobian and embedding perturbation of hidden representations. Layer-wise embeddings are further assessed for discrimination between homologous and non-homologous sequences, and model-predicted sequence likelihood differences between wild-type and mutant sequences are used for fitness prediction. (Lower right) Task-specific adaptation of LIMBER for downstream structural and functional inferences. The pre-trained model is fine-tuned and evaluated on secondary-structure prediction, sequence classification, IRES identification, MRL prediction and RNA–small molecule interaction prediction.

We further apply LIMBER to a broad panel of downstream tasks spanning secondary structure prediction, ncRNA classification, internal ribosome entry site (IRES) identification, mean ribosome load (MRL) prediction and RNA–small-molecule interaction prediction (Fig. 1, lower right), where LIMBER achieves consistently robust performance. Together, these results establish LIMBER not only as a high-performing RLM, but also demonstrate the application of a controlled platform to elucidate how architecture and scaling shape learned RNA representations. More broadly, the evaluation and interpretation framework provides a systematic basis for comparing RLMs, identifying transferable design principles to guide the future development of RLMs.

## 2 Results

### 2.1 Overview of LIMBER

LIMBER adopts a periodically interleaved Transformer–Transformer–Mamba (TTM) architecture (Fig. 1, left) designed to introduce complementary inductive biases for ncRNA sequence representation. The model is trained on *∼* 40.5 million ncRNA sequences from RNAcentral[9]. Additional details of the model architecture and pre-training procedure are provided in the Methods Sec. 4.1 and SI.

Self-attention captures direct couplings between arbitrary sequence positions and is naturally suited to capturing long-range nucleotide coevolutionary signals[10, 39]. Accordingly, most RLMs have adopted BERT architectures and achieved promising performance[17–20, 25–29, 40]. However, RNA sequences also exhibit intrinsically ordered organization that is not explicitly imposed by self-attention. RNA is a directional polymer synthesized in the 5*^′^ →* 3*^′^* direction[41, 42], and many recurring, conserved structural motifs display ordered signatures (e.g., tetraloops and C/D-box snoRNA motifs)[43–45], while thermodynamic models also capture sequence-asymmetric energetic contributions[46]. Mamba provides architectural priors for representing such ordered sequence features: its causal convolution integrates explicitly short-range sequential context, whereas the selective state-space mechanism adaptively updates contextual information along the sequence[11, 38]. Interleaving the two mechanisms therefore combines direct bidirectional global nucleotide interactions with local sequential context propagation. Given the finite and uneven sequence diversity represented in available datasets[9, 26], these complementary inductive biases could enable more effective learning of heterogeneous ncRNA sequence grammar.

To evaluate the hybrid architecture and characterize scaling behavior in RLMs, we train 10 LIMBER variants with differences in model size, pre-training-data scale, and causal convolution width, in addition to 3 variations of BERT models and 3 Mamba models. The three largest LIMBER variants contain approximately 100M, 300M and 650M parameters and are denoted LIMBER-*nano*, LIMBER-*meso* and LIMBER-*macro*, respectively. All pre-trained models are evaluated using the same task-agnostic evaluation and interpretation framework (Fig. 1, upper right), while representative LIMBER models are further adapted to various downstream tasks (Fig. 1, lower right).

### 2.2 Task-Agnostic Evaluation Framework Reveals Strong Intrinsic Capabilities of LIMBER

Masked-language-model pre-training requires models to infer masked nucleotides from sequence context, which encourages them to capture statistical dependencies within sequence space. We assess the biological relevance of these intrinsically learned dependencies are at three levels spanning structure, identity, and mutational fitness, enabling label-free, task-agnostic comparisons among LIMBER-*nano*,-*meso* and -*macro* and other RLMs. At the structural level, categorical Jacobian analysis[16] quantifies the response of model-output logits to all possible single-nucleotide substitutions and uses the resulting pairwise couplings to recover native contact maps. At the functional level, zero-shot classification evaluates whether learned representations distinguish homologous from non-homologous sequence pairs across RNA families and classes. Finally, zero-shot fitness prediction tests whether model-derived sequence variant likelihoods correlate with experimentally measured mutational effects, assessing whether learned sequence preferences reflect evolutionary constraints and biological activity. Full methodological details are provided in Methods Sec. 4.2–4.4. Together, these evaluations provide different views of the sequence-structure-function relationships encoded by RLMs during pre-training.

Applying the categorical Jacobian to the seven constituent datasets of bpRNA-1m(90)[47], comprising more than 25,000 RNAs across diverse families and types with structures derived from comparative analyses or experimental determination, we benchmark the LIMBER-family models against ten other models and two thermodynamic-based predictors using normalized area under the precision–recall curve (AUPR) metric (see Methods Sec. 4.2 for details on the datasets and evaluation metrics). Previous studies have attempted to directly reconstruct RNA contact maps from selected attention head, requiring the additional detection of informative heads and overlooking the feed-forward networks[27, 34]. Here, we instead apply the categorical Jacobian which provides a direct and end-to-end, full-network assessment. As shown in Fig. 2(a), performance varies significantly across RNA classes. LIMBER-*macro* consistently outperforms RNA-FM, structRFM, RNA-Ernie and LucaOne-family models[48] across nearly all RNA categories, and exceeds RNAstructure[49] and ViennaFold[50] for most cases, while showing broadly competitive performance with the RiNALMo-giga and AIDO-family models. At comparable model scales, LIMBER-*nano* also generally outperforms RiNALMo-mega. These results indicate that LIMBER effectively encodes covariational nucleotide signals required for the conservation of RNA secondary structures across a diverse sequence space. Moreover, increasing model scale generally improves structure recovery, a trend consistently observed across the LIMBER, RiNALMo and AIDO model families.

**Fig. 2.**
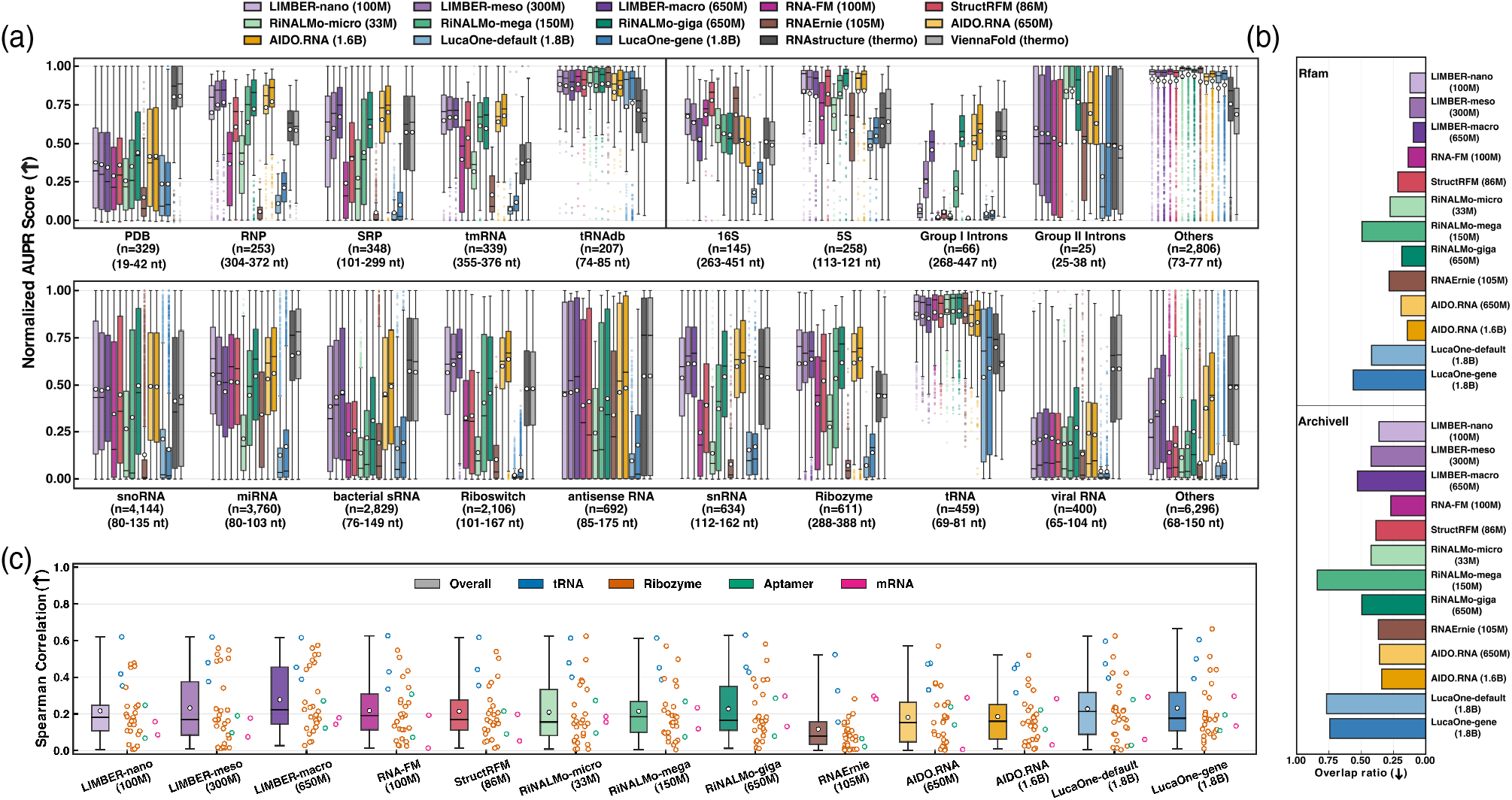
Task-agnostic Evaluation of LIMBER Models and Comparison with Other RNA Language Models. (a) Zero-shot evaluation of RNA secondary-structure recovery. Contact maps are directly predicted from each model using the Categorical Jacobian in the unsupervised manner, and performance is evaluated against annotated secondary structures using the Normalized AUPR metric. Seven datasets from bpRNA-1m(90) are evaluated: five are summarized in the upper-left panel, while CRW and Rfam are further stratified by RNA family in the upper-right and lower panels. For each dataset or RNA family, *n* denotes the number of sequences evaluated, and the first and third quartiles of the sequence-length distribution are indicated. Model sizes are specified in the legends. (b) Zero-shot evaluation of ncRNA classification. Overlap ratios between pairwise similarity distributions of homologous and non-homologous RNA sequences from Rfam (top) and ArchiveII (bottom) are computed from final-layer embeddings and compared across models. (c) Zero-shot evaluation of RNA fitness prediction. Absolute Spearman correlations between model-predicted likelihood scores and experimentally measured fitness or phenotypic effects are computed across 33 deep mutational scanning assays. Each assay is represented by a point, with colors denoting RNA types. The median (line) and mean (point) are indicated in each plot.

Furthermore, we assess the ability of RLMs to resolve functional relatedness by encoding sequence pairs drawn from the same or different RNA families (groups of evolutionarily related homologous sequences in Rfam 15.1 seed alignments[51]) and types (functional classifications in ArchiveII[52]), and quantifying pairwise similarity from their final-layer representations. The overlap ratio between the pairwise similarity distributions for homologous and non-homologous pairs provides a measure of discrimination, with lower overlap indicating better separation (see Methods Sec. 4.3). As shown in Fig. 2(b), LIMBER variants achieve leading family-level discrimination among all evaluated RLMs, while showing moderate, yet competitive, performance at the functional-type level. Across models, discrimination is consistently stronger on Rfam than on ArchiveII. Increasing model scale also improves family-level separation, whereas the opposite trend is observed for type classification. One plausible reason is that Rfam defines relatively narrow homology families with stronger conservation of sequence and structure patterns, which may align with the RLM pre-training objective. In contrast, broader ArchiveII RNA types can encompass much greater sequence divergence and weaker direct homology signals.

Additionally, we assess the ability of RLMs to predict the functional consequences of sequence variation using 33 deep mutational scanning assays from the RNAGym dataset, comprising 862,376 variants across ncRNAs and splicing-related RNAs[53]. These assays span diverse phenotypes, including RNA–protein binding for aptamers; self-cleavage, self-aminoacylation, template-directed RNA polymerization and self-splicing-dependent cellular fitness for ribozymes; cellular growth and translational suppressor activity for tRNAs; and pre-mRNA splicing efficiency for mRNA-based assays. Performance is quantified by the absolute Spearman correlation between experimentally measured phenotypic effects and the wild-type-conditioned log-likelihood ratios derived from RLM token logits (see Methods Sec. 4.4). LIMBER variants show relatively strong performance on ribozyme assays, with LIMBER-*macro* achieving the leading overall performance. Aptamer and splicing mRNA assays are generally more challenging across models, potentially reflecting differences in the representation of these sequence classes during pre-training. Increasing model scale also yields a clear performance improvement across the LIMBER family, whereas scaling trends are less evident for RiNALMo and AIDO models. We further benchmark RLMs across 23 additional coding mRNA assays[53] (Fig. S1). ncRNA-focused models generally perform less well on these tasks, including LIMBER, and unified protein–nucleic acid LucaOne models[48] show limited advantage. Overall, LIMBER models demonstrate strong and robust performance across three dimensions, and are directly comparable with many existing RLMs.

### 2.3 Scaling Behavior of LIMBER across Training, Model and Data

Understanding how training compute, model scale, and dataset size shape intrinsic RLM capabilities could inform more efficient model development. Given the model-size dependence observed across the unsupervised evaluations, we further systematically examine the scaling effects in LIMBER using the same benchmarks. Halving the training computes of the *nano*, *meso* and *macro* variants leaves classification and fitness-prediction performance largely unchanged, whereas secondary-structure recovery shows a slight dependence on training duration, particularly for longer RNAs (Extended Fig. 1). This trend parallels the continued reduction in model-predicted sequence perplexity (see definition in Method Sec. 4.1), showing that structural representations may benefit from additional pre-training compute.

Because extending training duration yields modest gains, we next examine scaling responses to model-size and dataset-size using model variants trained for half the original number of epochs to reduce computational cost. Increasing LIMBER capacity from 10M to 50M, 100M, 300M and 650M parameters produces broadly monotonic improvements in perplexity, secondary-structure recovery, family-level classification and fitness prediction, whereas RNA-type classification shows the opposite trend (Extended Fig. 2). Perplexity, structural recovery and family discrimination improve sharply from small to intermediate model scales, followed by progressively smaller gains at larger scales, whereas fitness prediction improves more steadily with model size. Similar size-dependent trends are also observed across the RiNALMo and AIDO model families (Fig. S3). Meanwhile, we examine data scaling by pretraining LIMBER-*nano* on 20%, 40%, 60%, 80% and 100% of the full training corpus (Extended Fig. 3). Perplexity and structural recovery show the largest gains up to approximately half of the full dataset, with additional data providing further benefits primarily for longer RNAs. Family classification and fitness prediction demonstrate more modest improvements with increasing data scale.

Consequently, under constrained computational budgets, increasing model capacity may provide larger marginal gains than extending training or enlarging the dataset once their moderate scales are reached. This diminishing return also motivates better data curation, for example, by improving sequence diversity rather than simply increasing corpus size. Classification capabilities appear comparatively easier to acquire, as even smaller models trained with less data and fewer epochs retain fair performance, whereas coevolutionary signals and prediction of mutational effects show stronger dependence on scaling. Improvements in latter capabilities coincide with weaker RNA-type discrimination, suggesting that distinct biological representations scale non-uniformly and may compete for shared model capacity.

### 2.4 Hybrid LIMBER Outperforms Pure BERT and Mamba Models

To date, newly developed RLMs have typically been benchmarked against earlier models that differ substantially in parameter count, training data and computational budget, confounding direct assessment of improvements arising from model design and training strategies, particularly given the scaling effects observed above. To isolate the contribution of the hybrid architecture, we apply the established unsupervised evaluation framework to pure BERT and Mamba models at 100M, 300M and 650M parameters, with all settings held identical to those of size-matched LIMBER models. Based on the training-scale behavior previously observed, all models are trained for half the number of epochs used for the original LIMBER models, reducing computational cost. A key motivation for incorporating Mamba is its 1-D causal convolution, which may capture short-range sequential motif grammar in ncRNA. To assess this contribution, we also train and test a LIMBER-*nano* variant with the convolution width reduced from 4 to 2, termed LIMBER-*nano-Conv2*.

Evaluation results for the four 100M, three 300M and three 650M models are summarized in Fig 3 and Figs. S4-S6. Across all model scales and evaluation tasks, LIMBER consistently outperforms the corresponding pure Mamba models. Relative to BERT, secondary-structure recovery based on categorical Jacobian analysis varies across RNA classes (Fig 3(a), Figs. S4(a) and S5(a)), while LIMBER achieves higher accuracies for 61%, 57% and 54% of the 26,707 test sequences at 100M, 300M and 650M parameters (Fig. S6), respectively. Using an embedding perturbation approach for unsupervised structure recovery (see details in Method Sec. 4.2 and following sections), these fractions are 72%, 76% and 53% (Fig. S6). Notably, LIMBER’s gains over BERT are enriched among RNAs longer than 100 nt. A similar length-dependent trend emerges when comparing LIMBER-*nano* with LIMBER-*nano-Conv2*: the longer convolution width outperforms the shorter variant for 57% of test sequences under either recovery approach, with the most differences concentrated among RNAs longer than 200 nt (Fig 3(a)).

**Fig. 3.**
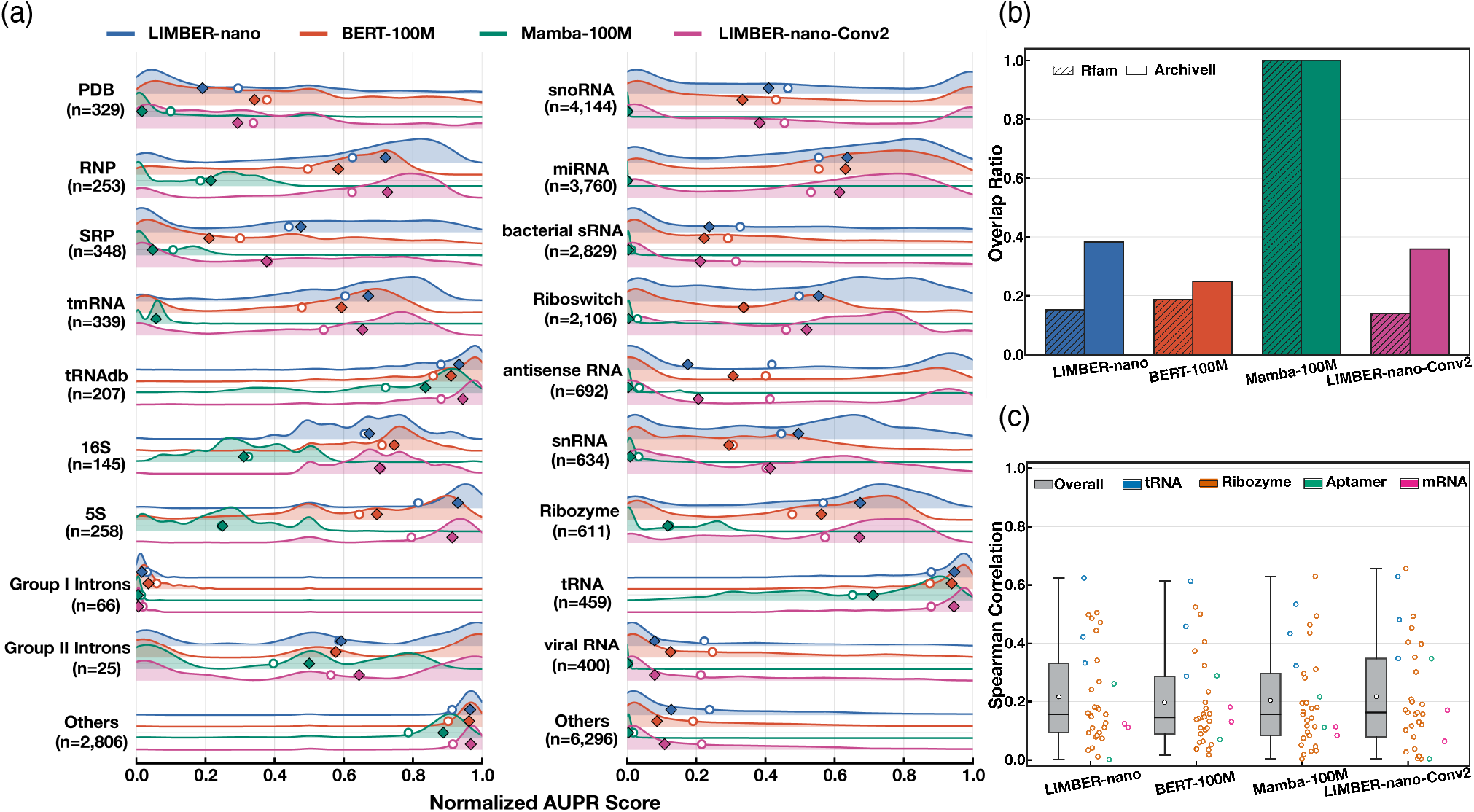
Hybrid LIMBER Architecture Improves Performance Over Pure BERT and Mamba Architectures. Four models with matched model size (*∼* 100M parameters) and training duration (i.e., 6 epochs, half the original schedule) are evaluated: LIMBER-*nano* (with a convolution kernel size of 4), BERT, Mamba, and LIMBER-*nano-Conv2* with a convolution kernel size of 2. (a) Unsupervised secondary-structure prediction across datasets using categorical Jacobian analysis, evaluated by normalized AUPR score. Mean and median scores are indicated by circles and diamonds, respectively. (b) Classification of homologous and non-homologous RNA sequences in model final-layer representations, quantified by the overlap ratio between similarity distributions. (c) Absolute Spearman correlations between model-derived relative likelihood scores and experimentally measured fitness across 33 mutational scanning assays.

RNA classification performance is less sensitive to model choice, with LIMBER and BERT variants showing broadly comparable results (Fig 3(b), Figs. S4(b) and S5(b)). LIMBER performs slightly better at the family level, whereas BERT shows a modest advantage at the RNA-type level, and convolution width has little effect on either task. Additionally, the hybrid architecture is observed to provide clear advantages for fitness prediction. While LIMBER and BERT yield comparable overall mean and median correlations across the 33 mutational assays at matched model scales, LIMBER models show clear gains for several ribozyme assays, yielding higher correlations in the upper quartile of the performance distribution (Fig 3(c), Figs. S4(c) and S5(c)). This advantage is retained when the causal convolution width is reduced. Overall, both cross-model comparisons (Sec. 2.2) and controlled architecture benchmarks indicate that the hybrid design balances diverse biological capabilities while maintaining effective representations of both RNA structure and function. Additionally, model-size scaling trends observed for LIMBER are also evident in BERT models (Fig. S7), suggesting that increasing model size provides general benefits across architectures.

### 2.5 Layer-wise Organization of Biological Information in LIMBER

Having established LIMBER’s robust performance across various evaluations, we further investigate the layer-wise organization of RNA biological knowledge inside the models. Two approaches are applied to characterize the contributions of individual layers and the progressive accumulation of biological information, respectively. Specifically, layer ablation strategy, which quantifies the effects of removing individual layers on model-output logits, combined with categorical Jacobian analysis and fitness likelihood scoring to assess layer-wise importance. Meanwhile, representations at successive layers are independently examined using the embedding perturbation method and zero-shot classification to characterize how structural and functional information emerges with model depth. Unlike categorical Jacobian analysis, which measures changes in logits induced by single-nucleotide substitutions, embedding perturbation[33] quantifies the corresponding changes in internal hidden representations at each model layer relative to the wild-type sequence (see Method Sec. 4.2). Secondary-structure recovery from final-layer embedding perturbation shows strong agreement with categorical Jacobian analysis across LIMBER model scales, with Pearson correlations exceeding 0.94 between their normalized AUPR scores (Fig. S2).

Layer-wise interpretation reveals consistent trends across LIMBER-*nano*,-*meso* and -*macro* models (Fig. 4 and S8). Ablation of Mamba blocks at layers 6, 9, and 30 of the encoder substantially impairs categorical-Jacobian-based secondary-structure recovery and weakens zero-shot fitness correlations, highlighting their critical roles in transforming representations. Ablation of individual Transformer layers has a slightly smaller impact on either task, while the final two Transformer layers exhibit greater contributions than earlier ones. Together with the controlled comparisons against pure BERT and Mamba models, these findings suggest that the hybrid architecture benefits from synergistic contributions of both layer types, particularly in the final few layers.

**Fig. 4.**
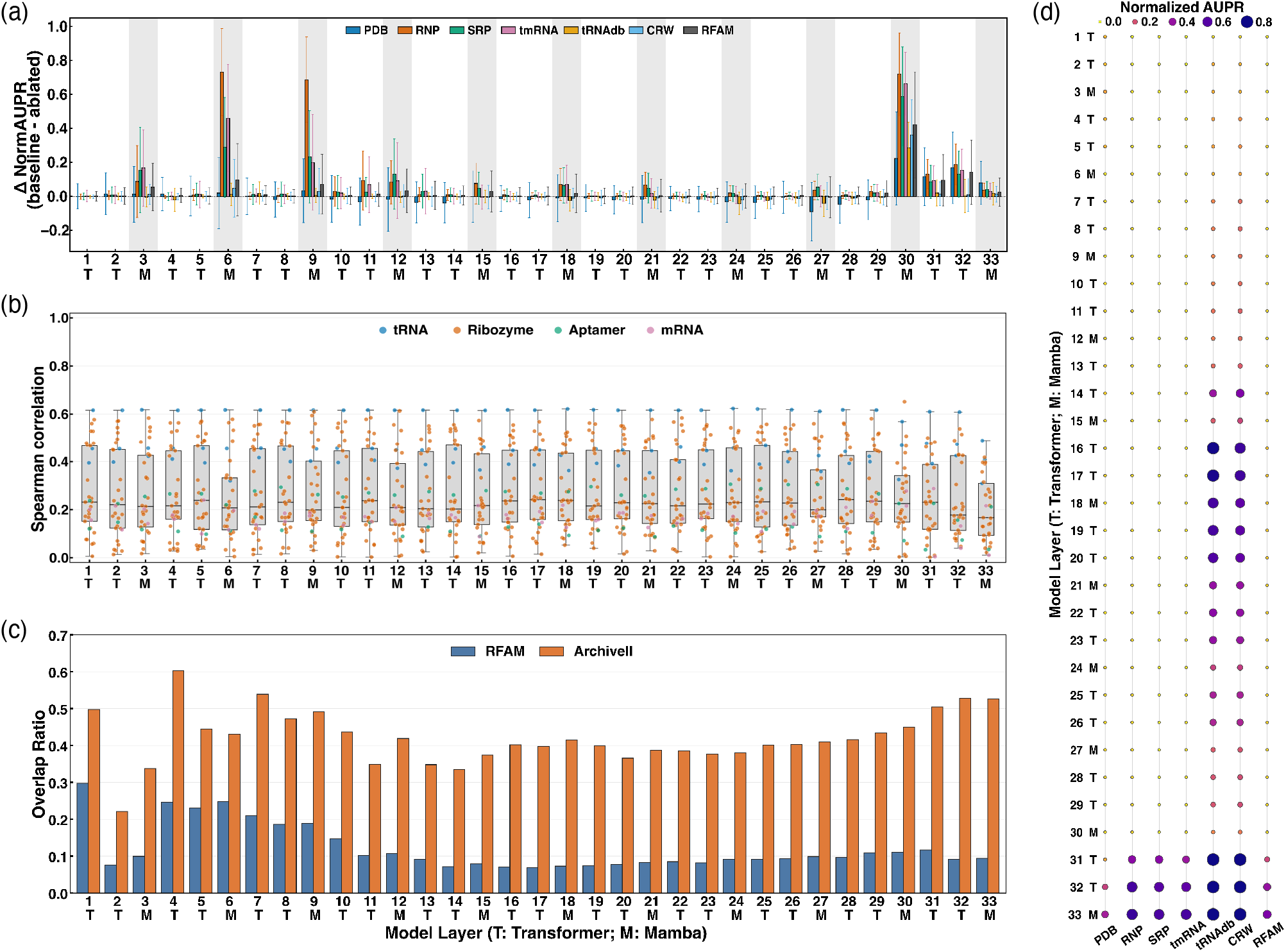
Layer-wise Interpretation of LIMBER Internal Representations for RNA Structure Recovery, Classification, and Fitness Prediction. Analyses are performed using the LIMBER-*macro* model trained for 12 epochs. (a) Effects of single-layer ablation on zero-shot secondary-structure recovery over seven bpRNA-1m(90) datasets, measured by changes in normalized AUPR from the categorical Jacobian method. (b) Effects of individual layer ablation on zero-shot fitness prediction across 33 deep mutational scanning assays, quantified by the absolute Spearman correlation between predicted likelihood score and experimental fitness for each layer-ablated model. (c) Layer-wise RNA family (Rfam) and type (ArchiveII) classification using embeddings extracted from successive model layers, quantified by the overlap ratio between pairwise sequence similarity distributions. (d) Progressive accumulation of RNA secondary-structure information across model layers, assessed using unsupervised embedding perturbation across seven datasets and quantified by normalized AUPR. Darker colors and larger markers indicate higher scores.

Embedding-perturbation analysis further reveals that structural information accumulates predominantly within the final two or three encoder layers. For some RNAs, structural signals emerge transiently at intermediate depths, weaken in subsequent layers and reappear near the output. Combined with the ablation results, these observations suggest that several intermediate layers contribute relatively little to output-logit predictions or structural recovery, potentially indicating redundant parameterizations. However, these seemingly uninformative representations at middle layers exhibit strong discrimination of RNA families and types, as reflected by lower overlap ratios in zero-shot classification. These findings reveal a depth-dependent, non-monotonic organization of biological information within LIMBER, in which structural recovery, logit likelihood prediction, and sequence discrimination exhibit different layer-wise dependencies. Therefore, deeper representations are not universally more informative and the utility of individual layers could depend on the biological task.

### 2.6 Fine-Tuned LIMBER Achieves Reliable RNA Secondary-Structure Prediction

Following conventional benchmarking protocols, we further assess LIMBER through supervised fine-tuning on various downstream prediction tasks, beginning with RNA secondary-structure prediction. Building on the coevolutionary information collected in LIMBER representations, we couple the pre-trained encoder to a pre-diction head that maps last-layer embeddings to pairwise base-pairing probabilities. The prediction head first constructs pairwise representations through a linear transformation of concatenated nucleotide pair embeddings, followed by two 2D convolutional blocks configured following the RiNALMo architecture[19]. Models are fine-tuned using a gradual unfreezing strategy with a binary cross-entropy loss; prediction accuracy and recall are quantified by F1 score. For fair comparisons, four baseline RLMs spanning *∼* 100M (RNA-FM and structRFM) and *∼* 650M (RiNALMo-giga and AIDO.RNA-650M) parameters are fine-tuned and evaluated under identical settings. Further details of the fine-tuning and evaluation procedures are provided in the Method Sec. 4.5 and SI.

We first evaluate RLMs using the curated bpRNA dataset following Ref. [54] in which sequences sharing more than 80% identity are filtered, yielding 13,419 RNAs partitioned into training (TR0, 10,814 sequences), validation (VL0, 1,300 sequences) and independent test (TS0, 1,305 sequences) sets. On TS0, all three LIMBER models outperform other baselines, with performance increasing from *nano*, *meso* to *macro*, consistent with the trends observed in unsupervised evaluations (Fig. 5(a)). To further assess model generalization to structurally distinct RNAs, we adopt the TrainSetA and TestSetB benchmarks introduced by Rivas et al.[55] TrainSetA comprises 2,943 sequences collected from published RNA structure benchmarks, whereas TestSetB contains 430 sequences from Rfam v10.0 seed alignments[56], with structures distinct from those in TrainSetA. Across this benchmark, all LIMBER variants outperform RiNALMo-giga, structRFM and RNA-FM, while AIDO.RNA-650M achieves performance intermediate between LIMBER-*macro* and LIMBER-*meso* (Fig. 5(b)).

**Fig. 5.**
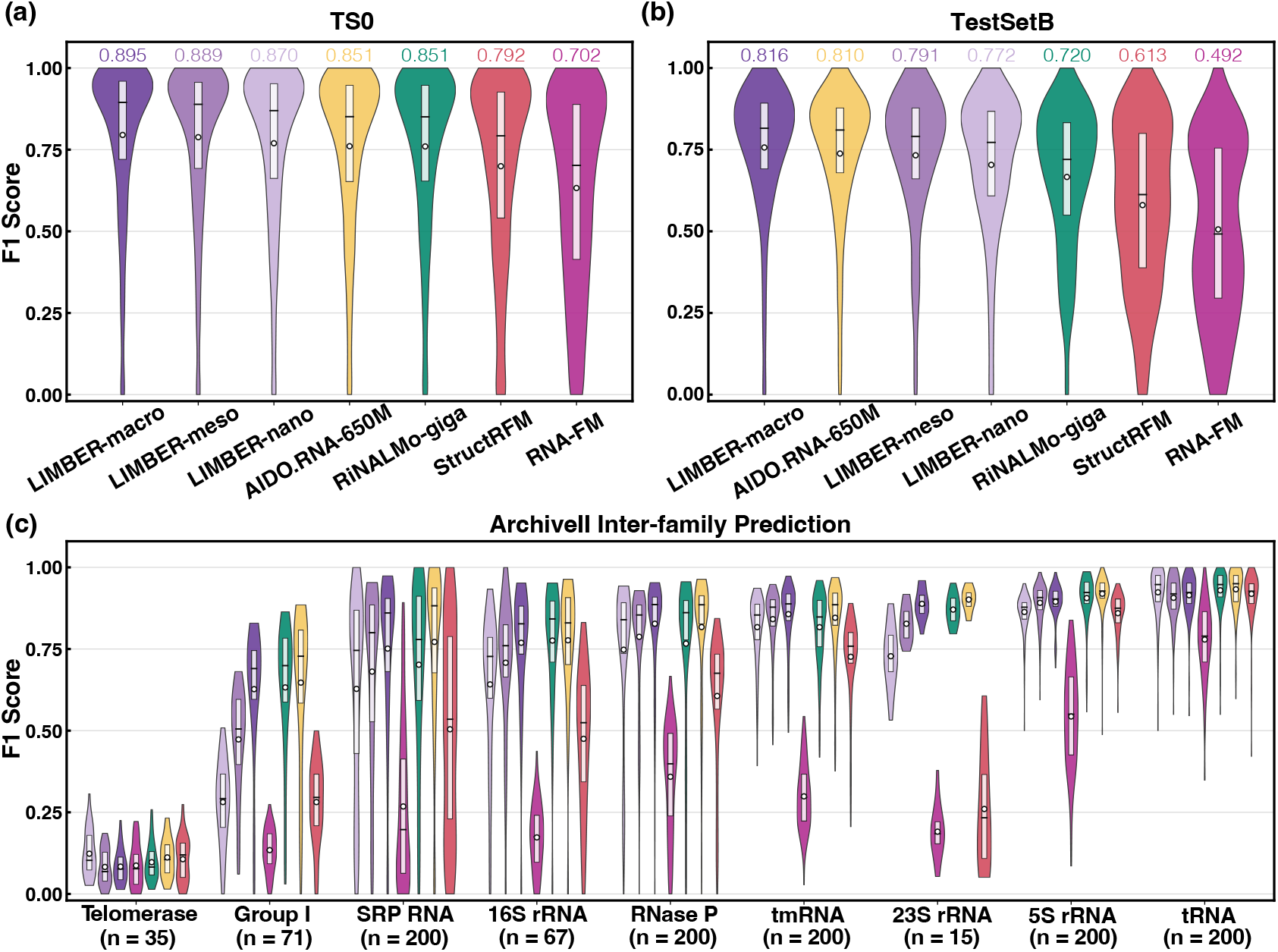
Fine-tuning LIMBER for RNA Secondary-Structure Prediction. LIMBER-nano,-meso and -macro are compared with four baseline models across secondary-structure prediction benchmarks using the flexible F1 score. (a) Models trained on TR0 and evaluated on TS0. (b) Models trained on TrainSetA and evaluated on TestSetB. In both panels, models are ranked by median F1 score, with its value annotated. (c) Cross-family evaluation on ArchiveII, with models trained on eight RNA families and evaluated on the held-out ninth family. The number of sequences in each held-out family is indicated. Across all panels, horizontal lines denote median scores, whereas points indicate means.

We next examine cross-family transferability using a leave-one-family-out evaluation on ArchiveII, following Ref. [57] and [19]. For each RLM, nine independent models are fine-tuned on eight RNA families and evaluated on the held-out ninth family. To reduce dataset imbalance, identical sequences are removed and each family is capped at 200 sequences, yielding 1,188 RNAs in total. As shown in Fig. 5(c), most models exhibit fair generalization across RNA families, with telomerase RNAs representing an exception. Telomerase RNAs are the longest among the nine families, though whether sequence length or pre-training data imbalance explains this behavior remains unclear. At comparable model scales, LIMBER-*nano* consistently outperforms RNA-FM and structRFM across all nine families, whereas LIMBER-*macro* performs comparably to AIDO.RNA-650M and RiNALMo-giga, achieving mean F1 scores of 0.809, 0.819, and 0.789, respectively. All benchmarks further confirm improvements in predictive performance with increasing model scale.

### 2.7 Fine-Tuned LIMBER Enables Accurate ncRNA Classification

We further assess LIMBER’s downstream adaptability through ncRNA classification. A classification head comprising dropout and a linear layer is applied to the final-layer [CLS] representation of LIMBER to predict RNA functional classes. Models are fine-tuned on the nRC benchmark[58], consisting of 6,320 sequences across 13 classes, and evaluated on a test set of 2,600 sequences, balanced at 200 per class. All other baseline RLMs are fine-tuned under identical settings. Additional details are provided in the Method Sec. 4.6 and SI.

As shown in Fig. 6(a), ncRNA classification is relatively tractable for pre-trained RLMs, with most models achieving strong predictive performance. LIMBER-*macro* leads across all five evaluation metrics–accuracy, area under the receiver operating characteristic curve (AUROC), area under the precision recall curve (AUPRC), Matthews correlation coefficient (MCC) and F1 score–while LIMBER-*nano* outperforms the size-matched RNA-FM and structRFM models. Noted by our earlier finding that functional class information is accumulated in middle layers (Fig. 4(c)), we additionally fine-tune a truncated LIMBER-*macro* model retaining only the first 14 encoder layers, denoted LIMBER-*macro (L14)*. Despite retaining only *∼* 45% of the original parameters, this variant achieves performance comparable to AIDO.RNA-650M and surpasses RiNALMo-giga, showing the potential of intermediate representations for efficient downstream adaptation. Class-specific F1 scores reveal broadly consistent performance patterns across models (Fig. 6(a), Right), with stronger predictions for well-represented RNA classes such as rRNA and tRNA, but weaker performance for miRNA, HACA-box and IRES elements. These differences may reflect imbalances in RNA-class representation within the pre-training data.

**Fig. 6.**
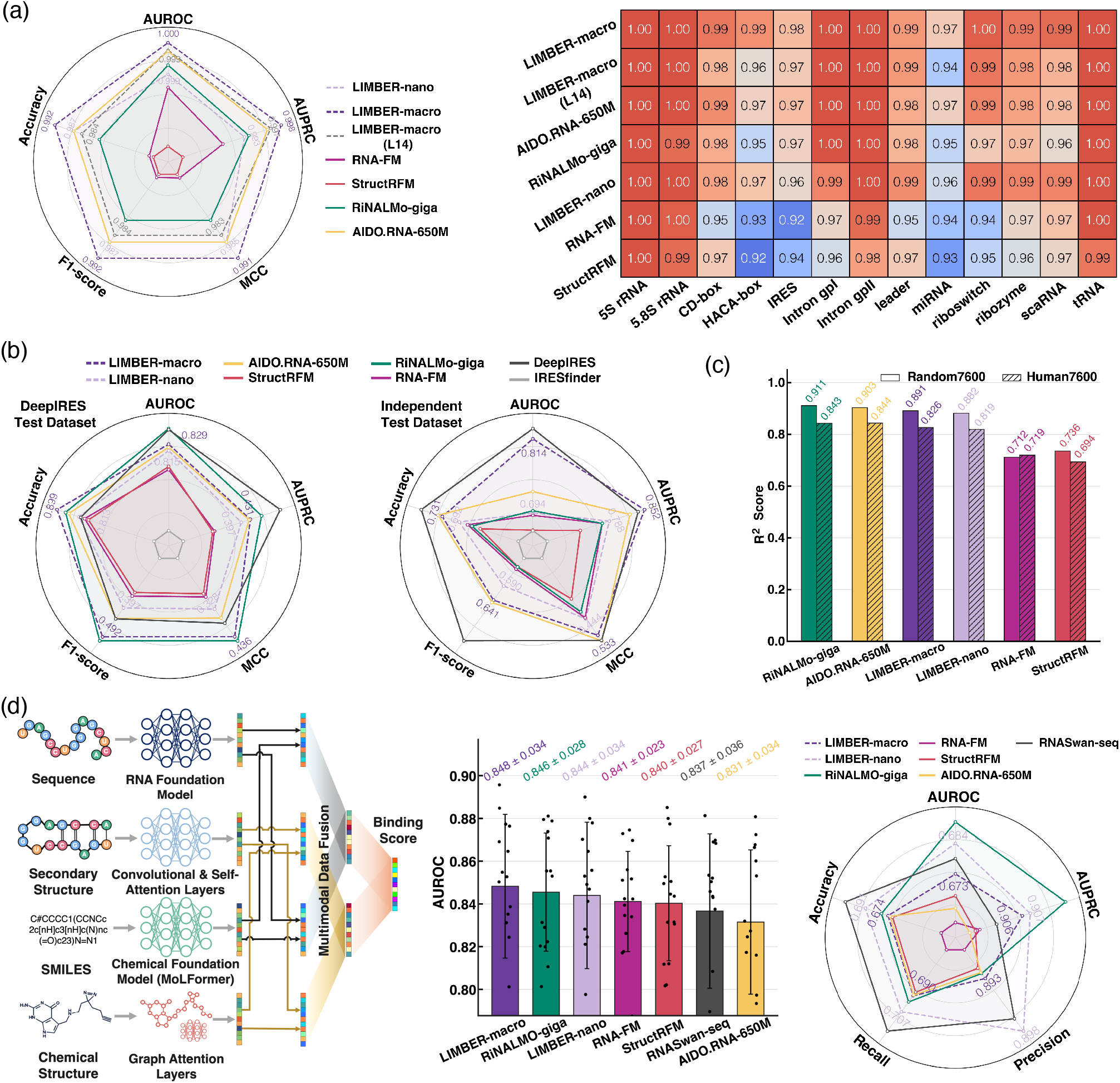
LIMBER-Driven ncRNA and IRES Classification, MRL Prediction and RNA–Small-Molecule Interaction Modeling. (a) *ncRNA Classification*; Left: Performance comparison of seven fine-tuned RLMs on the 13-class ncRNA classification task across five evaluation metrics. LIMBER models are visualized using dashed lines with annotated scores. Radial axes are independently shifted and linearly rescaled to improve visual separation between models. Right: Class-specific F1 scores of the seven models across 13 categories in the nRC dataset. (b) *IRES Identification*; Radar plots comparing IRES identification performance across five evaluation metrics for different RLMs and baseline methods on the DeepIRES test set (left) and an independent test set (right). Axes are independently shifted and linearly rescaled. (c) *MRL Prediction*; The squared Pearson correlation coefficient *R*^2^ between predicted and experimentally measured MRL values for the random and human 5*^′^* UTR test sets. (d) *RNA–Small-Molecule Interaction Prediction*; Left: Schematic of the multimodal prediction framework, adapted from the SMRTnet architecture. RNA sequence and secondary structure, together with small-molecule SMILES and chemical structure, are used as inputs to predict RNA–small-molecule binding scores. Middle: AUROC comparison of LIMBER-*nano*/macro and other RLMs on a fixed test dataset with a 1:2 positive-to-negative ratio. For each multimodal model, three independent training runs with different random seeds are performed using five-fold cross-validation, yielding 15 scores (black scatters); error bars indicate mean *±* s.d. Right: Performance comparison on SMRTnet benchmark datasets across five evaluation criteria. Values represent means over three runs with different random seeds. Radial axes are independently shifted and linearly rescaled, with results of LIMBER models highlighted.

### 2.8 LIMBER Captures Internal Ribosome Entry Site Activity and mRNA Translational Efficiency

Eukaryotic translation initiation is predominantly 5*^′^* cap-dependent, whereas IRESs enable cap-independent translation, particularly when canonical initiation is suppressed. IRES-mediated translation has been implicated in aberrant gene expression and associated with various diseases, highlighting its therapeutic relevance[59]. How-ever, experimental identification of IRES elements necessitates labor-intensive assays and controls, motivating efficient computational approaches for IRES prediction. Here, we couple the mean-pooled final-layer representations of a pre-trained LIMBER model with a feed-forward network that produces a single logit for classification of IRES and non-IRES sequences. The resulting model is fine-tuned and evaluated on the DeepIRES dataset[60], derived from high-throughput bicistronic assays, comprising 16,674 training and 1,853 test sequences, each 174 nt long, with an *∼* 1: 9 positive-to-negative ratio (see details on fine-tuning and baseline models in the Method Sec. 4.7).

On the DeepIRES test set, LIMBER-*macro* consistently outperforms AIDO.RNA-650M and the non-language-model baseline IRESfinder[61] across all evaluation metrics, while achieving performance comparable to RiNALMo-giga and the state-of-the-art task-specific method DeepIRES (Fig. 6(b), Left). We further test model generalization on an independently assembled benchmark comprising 52 IRES and 52 non-IRES sequences collected from the IRESfinder test set and previous experimental studies[60]. LIMBER-*macro* achieves the highest AUPRC, while performing slightly below DeepIRES on other metrics, comparably to AIDO.RNA-650M and much better than RiNALMo-giga (Fig. 6(b), Right). Across both benchmarks, LIMBER-*nano* consistently outperforms RNA-FM and structRFM models. These results demonstrate that LIMBER learns transferable sequence representations relevant to IRES activity.

The 5′ untranslated region (UTR) of mRNA plays a central role in regulating translation initiation by modulating ribosome recruitment and scanning[62]. Following initiation, multiple ribosomes can simultaneously translate the same mRNA, forming polysomes that support efficient protein synthesis. MRL, characterizes the average number of ribosomes associated with an mRNA molecule, serving as a proxy for translation efficiency[63]. Given the challenges of accurately predicting MRL from 5′ UTR sequences, we further evaluate the ability of LIMBER to capture sequence features governing ribosomal loading. Using a large-scale 5′ UTR random-library including 76,319 sequences paired with their MRL values[63], we fine-tune pre-trained RLMs by coupling their mean-pooled final-layer representations to a scalar regression head with the same architecture as the IRES classification head. The resulting models are assessed on two held-out test sets, each comprising 7,600 sequences, drawn from the same random library and from human 5′ UTRs, respectively (see Method Sec. 4.7 for details).

Using the squared Pearson correlation coefficient *R*^2^ as the evaluation metric, LIMBER-*macro* achieves values of 0.891 and 0.826 on random and human test sets, respectively, approaching the performance of RiNALMo-giga and AIDO.RNA-650M. And LIMBER-*nano* achieves similarly strong performance despite its smaller model size, substantially outperforming RNA-FM and structRFM models. Together with the IRES identification results, these findings show that LIMBER representations transfer effectively to mRNA regulatory tasks, despite being pre-trained predominantly on ncRNA sequences.

### 2.9 LIMBER Provides a Foundation for Modeling RNA–Ligand Interactions

Coding and non-coding RNAs play central roles in gene regulations and cellular processes, making RNA-dependent pathways attractive therapeutic targets. RNA-targeted small molecules have emerged as a growing focus in drug discovery, with the potential to modulate RNA functions or disrupt their interactions with other biomolecules[4, 5]. However, their computational design and screening remain limited by the scarcity of experimentally resolved structures, RNA intrinsic conformational flexibility, imperfect scoring functions for RNA–small-molecule interactions (RSMIs), and the under-exploration of chemical space for RNA-targeting small molecules[64, 65]. Given that LIMBER captures informative structural and functional features of RNA, we further investigate if it can facilitate RSMI prediction.

As illustrated in Fig. 6(d), we adopt the SMRTnet (predicting small molecule and RNA target interactions using deep neural networks) framework[66] to predict RSMI propensities by combining the RLM, the pretrained chemical language model MoLFormer[67], a convolutional and self-attention encoder, and a graph-attention encoder. Representations from the four networks are integrated through multimodal data fusion based on co-attention and self-attention, enabling joint modeling of cross-modal dependencies. The model processes RNA sequence and secondary structure together with small-molecule SMILES (simplified molecular-input line-entry system) and molecular structure to classify RSMIs in binary manner. SMRTnet incorporates its own Transformer-based RLM, RNASwan-seq, comprising 148M parameters and pre-trained on *∼* 214 million coding and non-coding RNA sequences. To isolate the contribution of the RNA representation, we replace RNASwan-seq with LIMBER variants and other RLMs while keeping the remaining SMRTnet architecture unchanged, and fine-tune each integrative model end-to-end on the original SMRTnet dataset. The dataset comprises 24,687 RNA–small-molecule pairs at a 1:2 positive-to-negative ratio, with experimentally derived secondary structures of the RNA binding fragments annotated. Training (80%), validation (10%) and test (10%) sets are partitioned such that no test-set SMILES appears in the training or validation data. Each multimodal model is trained in three independent runs with different random seeds using five-fold cross-validation, with predictions from the five folds ensembled for final scoring. Further details on dataset, model configuration, training procedures and scoring strategies are provided in the Methods Sec. 4.8 and SI.

LIMBER-driven SMRTnet models show robust discrimination between binding and non-binding RNA–small-molecule pairs, as measured by mean AUROC across five-fold cross-validation and three independent random seeds (Fig. 6(d)). LIMBER-*macro* surpasses RiNALMo-giga and AIDO.RNA-650M, whereas LIMBER-*nano* outperforms RNA-FM, StructRFM and RNASwan-seq. We further evaluate model transferability using another SMRTnet benchmark dataset comprising 2,011 experimentally characterized RSMIs, including 1,665 positive and 346 negative pairs curated from four datasets and 22 studies. Using the ensemble scoring strategy without prior knowledge of binding-pocket secondary structure (Method Sec. 4.8), both LIMBER variants maintain strong performance across five evaluation metrics (Fig. 6(d)), consistently outperforming RNA-FM, StructRFM and AIDO.RNA-650M, with LIMBER-*nano* slightly exceeding LIMBER-*macro*. Notably, SMRTnet benchmark data also includes RSMIs involving coding RNAs, showing that LIMBER representations may generalize beyond the pre-training data. These results support LIMBER as an effective foundation for RSMI prediction and RNA-targeted virtual screening and suggest that inclusion of explicit structural information about RNA binding sites is beneficial. In the future, incorporation of conformational ensembles for RNA and small molecules may further improve the physical fidelity and predictive performance[6, 68].

## 3 Conclusion & Discussion

Non-coding RNAs, which are more diverse and abundant than messenger RNAs, perform a wide range of bio-logical functions. Yet the scarcity of experimentally characterized structures and functional annotations limits predictive advances comparable to the success of AlphaFold in protein structural biology. The expanding repertoire of unlabeled ncRNA sequences offers another avenue: through self-supervised pre-training, RNA language models (RLMs) can learn patterns reflecting evolutionary, structural, and functional constraints, helping bridge RNA sequence and biological function.

Here, we introduce LIMBER, a hybrid RLM integrating bidirectional Transformer attention and Mamba selective state-space modeling. This architecture combines inductive biases for capturing long-range nucleotide dependencies and local ordered structural motifs. LIMBER demonstrates consistently strong performance and broad transferability across diverse downstream applications, including RNA secondary structure prediction, ncRNA classification, internal ribosome entry site (IRES) identification, mean ribosome load (MRL) prediction, and RNA–small-molecule interaction prediction. Beyond these benchmarks, we investigate a more fundamental question: what structural and functional information do RLMs intrinsically learn from RNA sequences? We develop a unified, task-agnostic evaluation and interpretation framework to probe the ability of pre-trained RLMs to recover secondary structures, distinguish sequence families and classes, and predict the effects of mutations on biological activity. These analyses reveal that LIMBER outperforms pure Transformer and Mamba architectures and organizes structural and functional information hierarchically across encoder layers, providing insight into the emergence of RNA representations during pre-training. Building on these findings, we systematically investigate the scaling of RLM capabilities with model size, pre-training dataset size and training compute. Within the regimes studied, model capacity emerges as a particularly influential factor in learning informative representations, providing ideas for the design of future RLMs.

Although LIMBER models have shown considerable promise, they remain insufficient to address many critical questions in RNA biology. We observe clear variation in structure and fitness prediction performance across RNA classes, potentially reflecting imbalances in existing sequence databases that limit representation learning and reduce training efficiency. Moreover, the competitive performance of conventional computational biophysics methods on certain tasks suggests that important thermodynamic and physical constraints may not be fully captured by sequence-only pre-training approaches. Incorporating complementary structural inductive biases, such as structure-informed self-attention in ERNIE-RNA[26] and structure-guided masking in structRFM[27], together with more systematic dataset curation and sampling strategies, may yield further improvements. More fundamentally, a comprehensive understanding of RNA function and rational RNA-targeted drug discovery requires characterization of heterogeneous conformational ensembles and their dynamics, rather than single static structures[6, 68]. However, high-quality data resolving these ensembles remains limited, and their generation through experiments or molecular dynamics simulations is technically and computationally demanding. This limitation motivates an intriguing future direction: steering the evolutionary and structural constraints learned by RLMs to generate diverse, physically plausible RNA conformations beyond a single predicted structure.

Finally, while increasing model capacity and architectural complexity may yield further improvements, we emphasize the importance of closing the scientific loop between model development, rigorous evaluation, and mechanistic interpretation. Systematic benchmarking across complementary dimensions, together with a deeper understanding of what models learn, how learned information is organized within and across layers, and where the models fail, can provide a more principled basis for developing the next generation of RLMs.

## 4 Methods

### 4.1 LIMBER Model Architecture and Pre-training

#### Model configurations

LIMBER follows a repeating TTM pattern, in which Transformer layers use multi-head padding-aware self-attention and position-wise feed-forward networks, whereas Mamba layers operate in a pre-normalized residual formulation with a 16-dimensional state space, a causal convolution width of 4, and an expansion factor of 4. Input RNA sequences are tokenized at single-nucleotide resolution and represented using learned nucleotide and absolute positional embeddings. The final hidden embeddings are passed to a masked-language-modeling head and projection to the nucleotide vocabulary. The vocabulary contains five special tokens (i.e., [PAD], [UNK], [CLS], [SEP], and [MASK]), four canonical RNA nucleotides (i.e., A, U, C, and G), and an ambiguous nucleotide token N. Sequence preprocessing converts all nucleotides to uppercase, replaces thymine with uracil, and maps non-ACGU symbols to N.

#### Pre-training Procedure

Sequences from RNAcentral[9] are used for pre-training after pre-processing and exact-sequence de-duplication, yielding 40,498,819 RNA sequences together with a fixed validation set. Sequences *≤* 1,022 nt are used in full, whereas longer sequences are randomly cropped to 1,022 nt. Models are pre-trained using masked language modeling, where 15% of nucleotide positions are selected for prediction. Of these, 80% are replaced by [MASK], 10% by a randomly sampled canonical nucleotide, and 10% are left unchanged; cross-entropy loss is computed only over the selected positions. LIMBER is instantiated at multiple model scales while preserving the same TTM architecture. Detailed architectural configurations, pre-training data, and optimization settings are provided in the SI.

#### Perplexity Calculation

Perplexity measures the predictive fidelity of a language model’s predictive fidelity. To avoid the computational cost of evaluating full-sequence likelihoods, we estimate perplexity from the average negative log-likelihood of selected nucleotide positions:

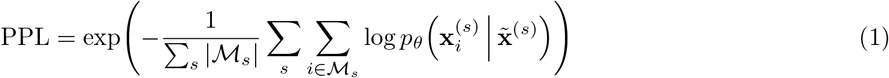

where **x**^(*s*)^ is the *s*-th validation sequence, *M_s_* is the set of masked positions in the sequence, and **x̃**^(*s*)^ is the corrupted input. Approximately 15% of nucleotide positions are randomly selected and corrupted following the masked language modeling procedure described above.

### 4.2 Unsupervised Secondary Structure Recovery: Categorical Jacobian and Embedding Perturbation

Zero-shot evaluation provides a stringent test of secondary-structure prediction, assessing whether the model has learned transferable coevolutionary signals rather than dataset-specific patterns introduced through supervised fine-tuning. We employ two approaches, the categorical Jacobian[16] and embedding perturbation[33], to directly extract native secondary-structure contact maps from the pre-trained model in a fully unsupervised manner.

#### Categorical Jacobian

Given an RNA sequence *X* = *{x*_1_*, x*_2_*, · · ·, x_L_}* of length *L* over an alphabet of A nucleotide tokens (A=4, with only A, U, C, and G considered here), the categorical Jacobian **J**[16] is an tensor with size of *L × A × L × A* that quantifies the sensitivity of language-model output logits to all possible single-nucleotide perturbations of the input sequence. To construct **J**, each nucleotide position is substituted with every possible token, yielding *L × A* sequence variants, each of which is further passed through the language model *P* to obtain an *L × A* logit matrix. Differences between these logits and those obtained for the original sequence **J***_ij_* = *P*[*X* (*x_i_ → A_j_*)]^(L×A)^ *− P*[*X* ^](L×A)^ are then assembled into the categorical Jacobian. After mean-centering **J** along each dimension, the two alphabet dimensions are collapsed using the Frobenius norm to yield an *L × L* matrix *C_ij_* = *|***J**[*i,*:, *j*,:]*|*^2^. This matrix is subsequently symmetrized *C_ij_ ←* (*C_ij_* + *C_ji_*)*/*2 followed by Average Product Correction (APC)[69] to reduce background noise and phylogenetic bias, 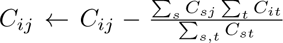, yielding the final contact map matrix.

#### Embedding Perturbation

Whereas the categorical Jacobian captures output-logit responses to token perturbations, embedding perturbation probes the corresponding changes in RLM layer-wise internal representations[33]. Similarly, *L × A* sequence variants are generated by substituting each nucleotide at every sequence position and are individually passed through the language model. At layer *k*, the hidden representation **z**^(*k*)^ *∈ ℝ*^L×c^ for each sequence is normalized across each hidden channel (where *c* denotes the hidden channel dimension), and its deviation from the wild-type representation is used to construct a sensitivity tensor of size *L × A × L × c* over all sequence variants. As above, the alphabet and hidden-channel dimensions of the tensor are collapsed using the Frobenius norm, followed by symmetrization and APC[69]. This procedure yields a contact map from each model layer.

#### Normalized Area Under the Precision-Recall Curve (AUPR) Score

Ground-truth secondary structures in dot-bracket notation are converted to binary contact matrices, whereas unsupervised methods produce continuous contact scores without a fixed scale. Performance is therefore quantified using normalized AUPR. We consider only entries in the upper triangle of the predicted contact matrix and exclude nucleotide pairs separated by fewer than four positions along the sequence. Predicted contact scores are paired with the corresponding ground-truth binary labels and ranked to construct the precision–recall curve, with AUPR obtained by trapezoidal integration. AUPR is further normalized against the expected performance of a random predictor, such that 1 denotes perfect prediction and 0 denotes random performance[33, 39]. For the two thermodynamic predictors, RNAstructure[49] and ViennaRNA[50], base-pairing probability matrices are used for evaluation. Predictions are performed at 60*^◦^*C, following prior studies[39, 70], with all other parameters set to their default values.

#### Secondary Structure Dataset

The integrative bpRNA-1m(90) dataset[47] is used for evaluation. The dataset aggregates RNA sequences and annotated secondary structures from the RCSB Protein Data Bank (PDB), Nucleic Acid Database, Signal Recognition Particle (SRP) Database, tmRNA Database, tRNAdb 2009, Comparative RNA Web (CRW), and all families in Rfam 12.2. To reduce redundancy, sequences with *>* 90% identity over at least 70% alignment coverage were removed. After further excluding sequences longer than 512 nucleotides, the final benchmark contains 26,707 sequences, with source-specific counts summarized in Fig. 2. Additional details of the dataset are provided in the SI.

### 4.3 Zero-shot RNA Classification

The language model’s ability to capture functional similarity is evaluated using two classification benchmarks at the family and type levels. Homologous and non-homologous sequence pairs are constructed independently from Rfam 15.1 seed alignments[51] and ArchiveII[52], respectively, with 100,000 pairs sampled for each class.

#### Dataset Construction

After restricting sequence length to 512 nt and removing alignment gaps, the Rfam 15.1 seed alignments[51] yield 108,256 sequences across 4,194 families, whereas ArchiveII[52] dataset contains 3,865 sequences spanning nine broader RNA classes after exclusion of the long group II intron class. Homologous and non-homologous pairs are sampled from the same and different families or RNA classes, respectively. Sampling is performed uniformly and at random without replacement, with self-pairs excluded, and each unordered pair is retained at most once. It is noted that 511 sequences occur more than once in ArchiveII, and this redundancy is retained during sequence-pair construction.

#### Sequence Similarity

RNA sequence pairs are encoded by the trained model, and hidden representations are extracted after each layer and transformed into fixed-length spectral features using the fast Fourier transform (FFT). The real components of the FFT coefficients are retained, and their cosine similarity is used to quantify the similarity of RNA-pair representations across model layers[34]. Class discrimination is scored as the overlap ratio between the resulting similarity distributions of homologous and non-homologous pairs, where lower overlap of homologous and non-homologous distributions indicates better class identification.

### 4.4 Zero-shot Fitness Prediction

Given a wild-type sequence *X_wt_* = *{x*_1_*, x*_2_*, · · ·, x_L_}*, the likelihood score of a mutant sequence *X_mut_* carrying mutations at positions *M ⊆ {*1, 2*, · · ·, L}* is computed as[34, 71]:

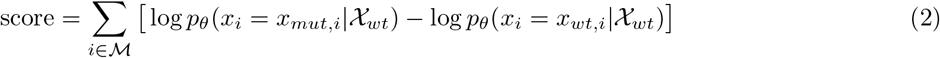

where log *p_θ_*(*x_i_|X*) denotes the model-assigned probability of nucleotide *x_i_* at position *i*, conditioned on sequence context *X*. The applied wild-type-conditioned log-likelihood ratio quantifies the model’s relative support for each mutant nucleotide compared with its wild-type counterpart within the same sequence context. Since all mutation scores are derived from logit predictions conditioned on *X_wt_*, the calculation requires only a single forward pass of the wild-type sequence through the model.

### 4.5 Fine-tuning LIMBER for Secondary Structure Prediction

#### Prediction Head Architecture

Given final-layer embeddings from an RLM of a sequence **z** *∈ ℝ*^L×c^, pairwise representations are constructed as **p***_ij_* = [**z**[*i*]; **z**[*j*]] and linearly projected to 64 dimensions. The resulting *L×L×* 64 tensor is processed by two convolutional residual blocks, followed by a final convolution layer that produces an *L × L* matrix of base-pairing logits. The matrix is then symmetrized, and binary cross-entropy loss is computed over its strictly upper-triangular logits.

#### Fine-tuning and Structure Decoding

A gradual unfreezing strategy is adopted, with only the prediction head optimized during the first three epochs. Subsequently, three additional encoder layers are unfrozen every three epochs, proceeding from the final layer toward the input and fine-tuned at a smaller learning rate. During inference, base pairs are assigned through greedy matching using a validation-tuned threshold, subject to canonical and wobble pairing constraints and the exclusion of short hairpin loops. Performance is evaluated using a flexible F1 score, which permits a one-nucleotide positional mismatch in predicted base pairs, following prior studies[19, 72]. All baseline models are fine-tuned using identical configurations and evaluation protocols. Further details on model architecture, training, and evaluation are provided in the SI.

### 4.6 Fine-tuning LIMBER for ncRNA Classification

The [CLS] representation is passed through layer normalization, dropout (rate 0.1), and a linear classifier to predict probabilities across 13 RNA classes. Fine-tuning proceeds in two stages using cross-entropy loss: the RLM backbone is frozen while the classification head is trained for 10 epochs, followed by end-to-end fine-tuning for an additional 20 epochs at a reduced learning rate. Class predictions are assigned by the maximum predicted probability; AUROC and AUPRC are computed using a one-versus-rest strategy across all 13 classes.

### 4.7 Fine-tuning LIMBER for IRES Identification and MRL Prediction

#### IRES Identification

For each input sequence, the RLM representations are mean-pooled across nucleotide positions, excluding [CLS], [SEP] and padding tokens. The pooled representation is passed through layer normalization, dropout (rate 0.1), and a two-layer feed-forward network for binary classification. Fine-tuning proceeds in two stages with cross-entropy loss. First, the pre-trained RLM backbone is frozen, and only the classification head is optimized for 8 epochs, retaining the checkpoint with the highest validation AUROC. The backbone is then unfrozen, and the entire model is fine-tuned for an additional 12 epochs at a reduced learning rate. The checkpoint achieving the highest validation AUROC is selected for inference. A validation set comprising 10% of the 16,674 training sequences is randomly held out. A fixed decision threshold of 0.5 is used to calculate threshold-dependent evaluation metrics.

#### MRL Prediction

The MRL prediction head follows the same architecture and fine-tuning strategy as the IRES classifier head, using mean-pooled final-layer RLM representations to predict a scalar MRL value. Training follows the same two-stage protocol using a Huber regression loss. The checkpoint with the best validation performance is selected using a randomly held-out subset comprising 10% of the training data. All baseline models are fine-tuned and evaluated under identical configurations. Further details on IRES identification and MRL prediction are provided in the SI.

### 4.8 LIMBER-Driven Prediction of RNA–Small-Molecule Interactions

#### Model Configurations

We adopt the SMRTnet architecture[66], replacing RNASwan-seq with each RLM variant while retaining MoLFormer[67] as the chemical language model. RLM representations with hidden dimensions other than 640 are projected to 640 dimensions through a linear layer. The RNA secondary-structure encoder, molecular graph-attention encoder, and multimodal fusion module use the original SMRTnet architectures[66] and are randomly initialized. Each integrative model is optimized end-to-end with a batch size of 32 for up to 100 epochs, using binary cross-entropy loss with *L*_2_ regularization and early stopping based on validation AUROC (patience 20 epochs). Each multimodal model is independently initialized and trained with three random seeds (i.e., 1, 2 and 42) across five-fold cross-validation, yielding 15 trained models in total. For benchmarking RNASwan-seq, we do not directly use the public SMRTnet weights; instead, consistent with the evaluation of other RLMs, the two structural encoders are randomly initialized and trained from scratch.

#### Dataset Construction

For training, validation and testing, we employ exactly the identical five-fold cross-validation splits of the SMRTnet dataset as reported in [66]. The dataset is derived from 1,061 selected RNA–small-molecule complex structures in the PDB, with experimentally resolved interactions serving as positive pairs and negative pairs generated by random RNA–small-molecule pairing after excluding known interactions, yielding 24,687 pairs at a 1:2 positive-to-negative ratio. Following the original SMRTnet protocol, only 31-nt RNA fragments centered around the binding pocket, together with their corresponding secondary structures, are used as input for training. Additional validation is performed on the SMRTnet benchmark dataset assembled from four resources (R-BIND, R-SIM, SMMRNA and NALDB) and 22 publications[66], after removing RSMIs overlapping with the SMRTnet dataset.

#### Ensemble Scoring Strategy and Model Evaluations

Following [66], experimentally derived secondary structures of 31-nt RNA binding-site fragments are provided as inputs during both training and testing of the multimodal models, and test-set performance is evaluated by AUROC for each of the 15 trained models. For the SMRTnet benchmark dataset, we adopt an ensemble scoring procedure. Each RNA is scanned using 31-nt sliding windows with a stride of 1 nt, and every window, together with its corresponding experimentally resolved secondary structure, is evaluated by the five cross-validation models. The median of the five predicted probabilities is taken as the window-level binding score. For RNAs of 31–40 nt, the maximum score across all windows is used as the final score. For RNAs longer than 40 nt, candidate binding regions are identified as four consecutive windows with score *>* 0.5, and the maximum score among four windows is used as the final score. If no such region is detected, the minimum score across all windows is assigned, indicating a non-binding interaction. Performance is quantified by accuracy, precision, recall, AUPRC and AUROC, with each metric averaged across the three independent training seeds. Although this strategy does not require prior binding-site localization, its local-window design may underrepresent long-range RNA interactions. More details are presented in the SI.

## 5 Data & Code Availability

All code for pre-training, evaluation and downstream adaptation is available at https://github.com/ziyaow1010/LIMBER. Pre-trained weights for every LIMBER variant, together with the size-matched pure-BERT and pure-Mamba models and the data- and compute-scaling checkpoints, are released at https://huggingface.co/Ziyao1010/LIMBER, one subfolder per model. The RNA sequences used for pre-training are obtained from the RNAcentral database (https://ftp.ebi.ac.uk/pub/databases/RNAcentral/currentrelease/sequences/). For RNA secondary structure evaluation, we use bpRNA-1m (https://bprna.cgrb.oregonstate.edu/), the SPOT-RNA TR0/VL0/TS0 splits of bpRNA (https://github.com/jaswindersingh2/SPOT-RNA), the Rivas datasets distributed with MXfold2 (https://github.com/mxfold/mxfold2/releases), and the ArchiveII family-based splits (https://github.com/marcellszi/dl-rna). Classification benchmarks include Rfam (https://rfam.org/), ArchiveII, and the 13-class nRC dataset (https://github.com/IcarPA-TBlab/nrc). For unsupervised mutational fitness prediction, we used deep mutational scanning dataset from RNAGym (https://huggingface.co/datasets/Marks-lab/RNAgym). Functional evaluation datasets comprise the DeepIRES benchmark (https://github.com/SongLab-at-NUAA/DeepIRES); the variable-length 5*^′^* UTR massively parallel reporter assay dataset (GEO accession GSE114002; https://www.ncbi.nlm.nih.gov/geo/query/acc.cgi?acc=GSE114002); and the SMRTnet training and benchmark datasets for RNA–small-molecule interaction prediction (https://zenodo.org/records/14715564).

## 6 Acknowledgment

This work is partially supported by NSF Award (2607564). Y.Q. acknowledges support from startup funds provided by the University of Notre Dame. Computational resources are provided by the the University of Notre Dame Center for Research Computing and the University of Maryland Institute for Health Computing. The authors acknowledge the use of AI tools, including Claude and ChatGPT, to assist with code development and grammatical editing of the manuscript.

---

**Extended Data Fig. 1.**
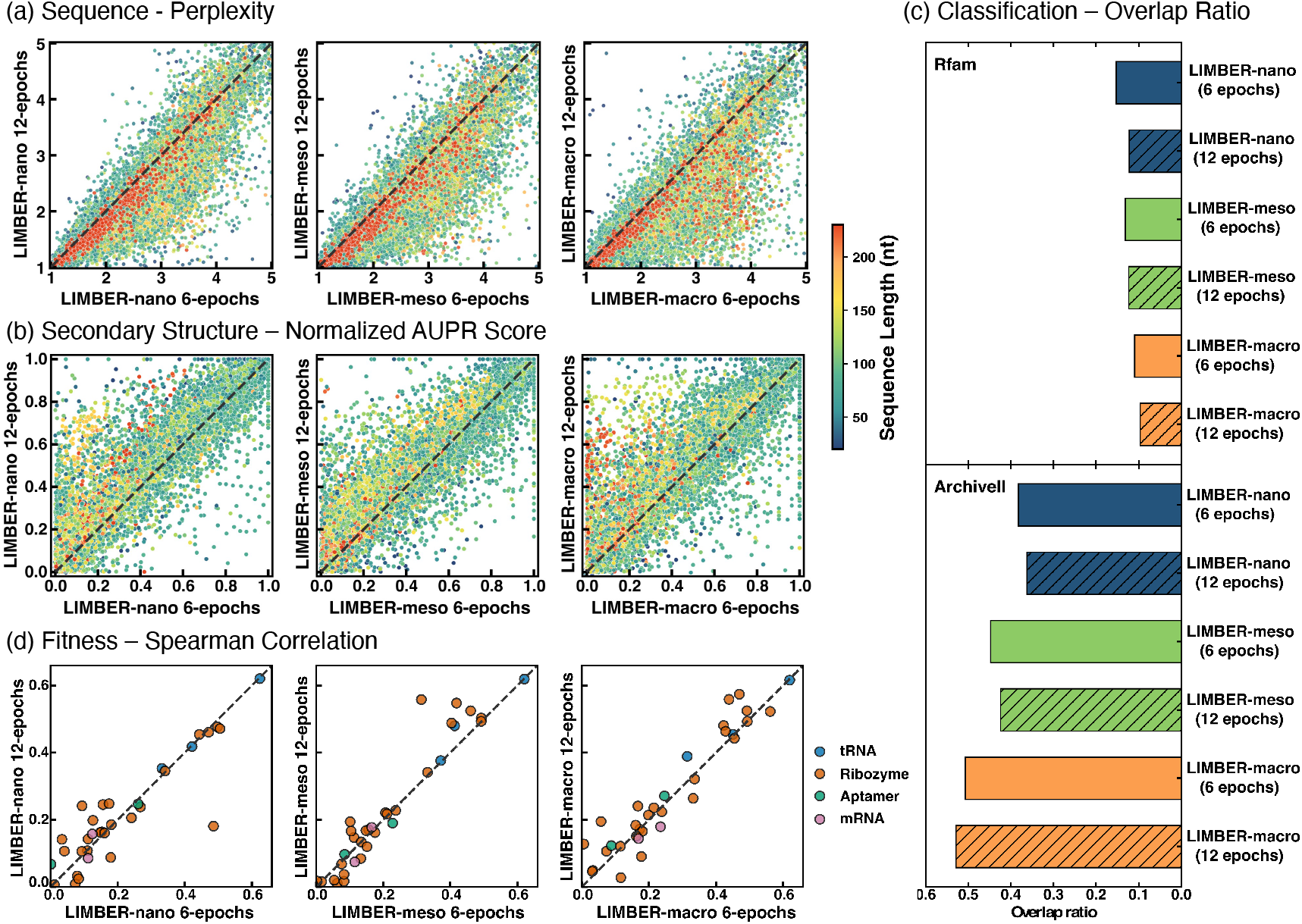
Emergence of RNA structural and functional representations with increasing training compute. LIMBER-*nano*, LIMBER-*meso*, and LIMBER-*macro* are trained on the full dataset for six or twelve epochs. (a) Sequence-level perplexity across the cleaned bpRNA-1m(90) dataset. (b) Zero-shot secondary-structure prediction performance on bpRNA-1m(90), quantified by normalized AUPR using the categorical Jacobian method. In (a,b), each point represents one RNA sequence and is colored by sequence length. (c) Zero-shot RNA family classification across LIMBER models trained for different numbers of epochs, evaluated by the overlap ratio between pairwise similarity distributions from final-layer embeddings of homologous and non-homologous sequences sampled from the Rfam and ArchiveII datasets. (d) Zero-shot RNA fitness prediction across 33 deep mutational scanning assays, quantified by absolute Spearman correlation between model-predicted likelihood scores and experimentally measured fitness.

**Extended Data Fig. 2.**
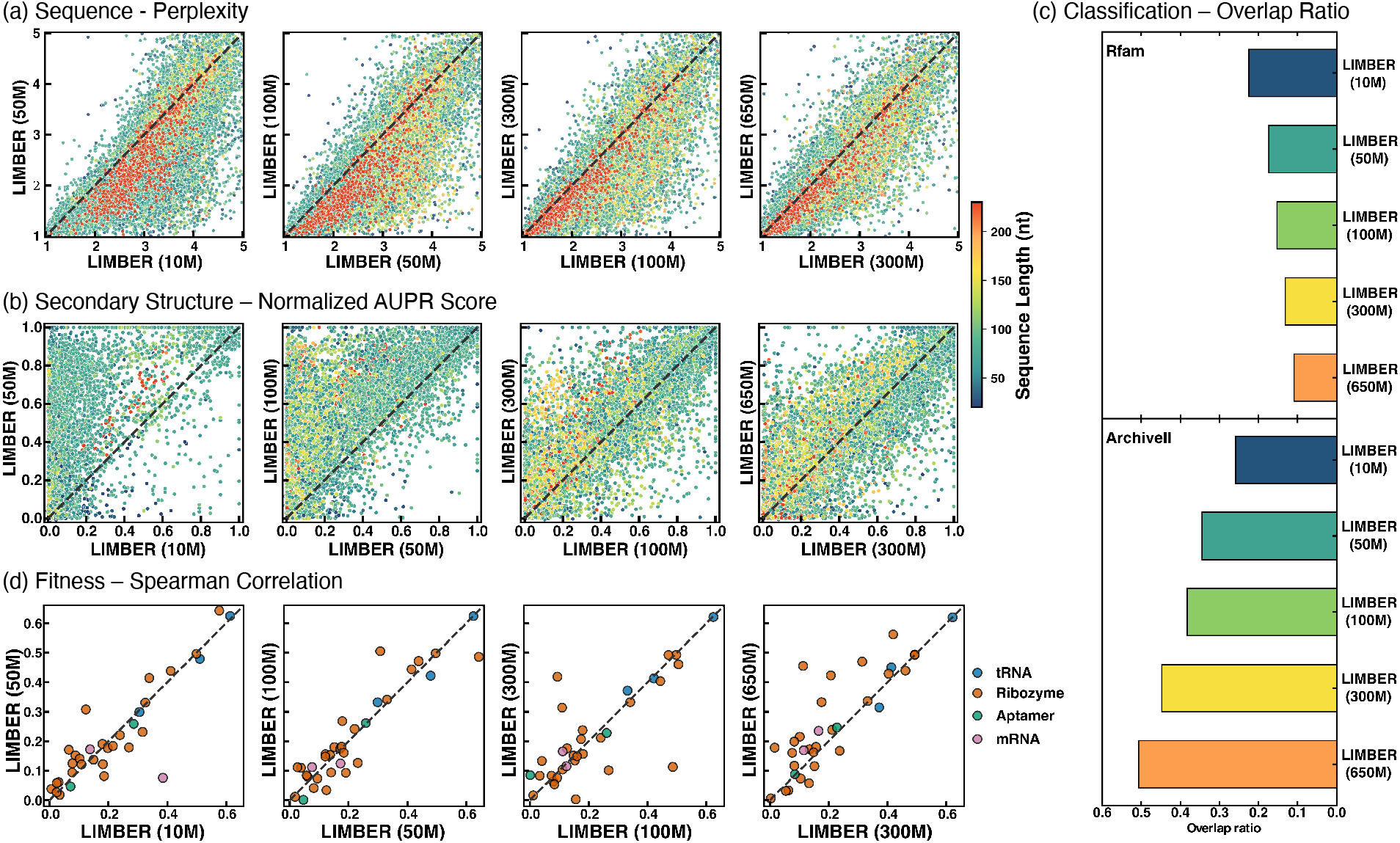
Emergence of RNA structural and functional representations with LIMBER model scaling. LIMBER models of increasing parameter size are trained on the full dataset for 6 epochs. Performance is evaluated using the same perplexity, secondary-structure, and fitness benchmarks described in Extended Fig. 1. (a) Sequence-level perplexity for each model compared with the next larger model across bpRNA-1m(90). (b) Zero-shot secondary-structure prediction performance. (c) Zero-shot RNA classification for homologous and non-homologous sequences sampled from the Rfam and ArchiveII datasets. (d) Zero-shot RNA fitness prediction across 33 deep mutational scanning assays.

**Extended Data Fig. 3.**
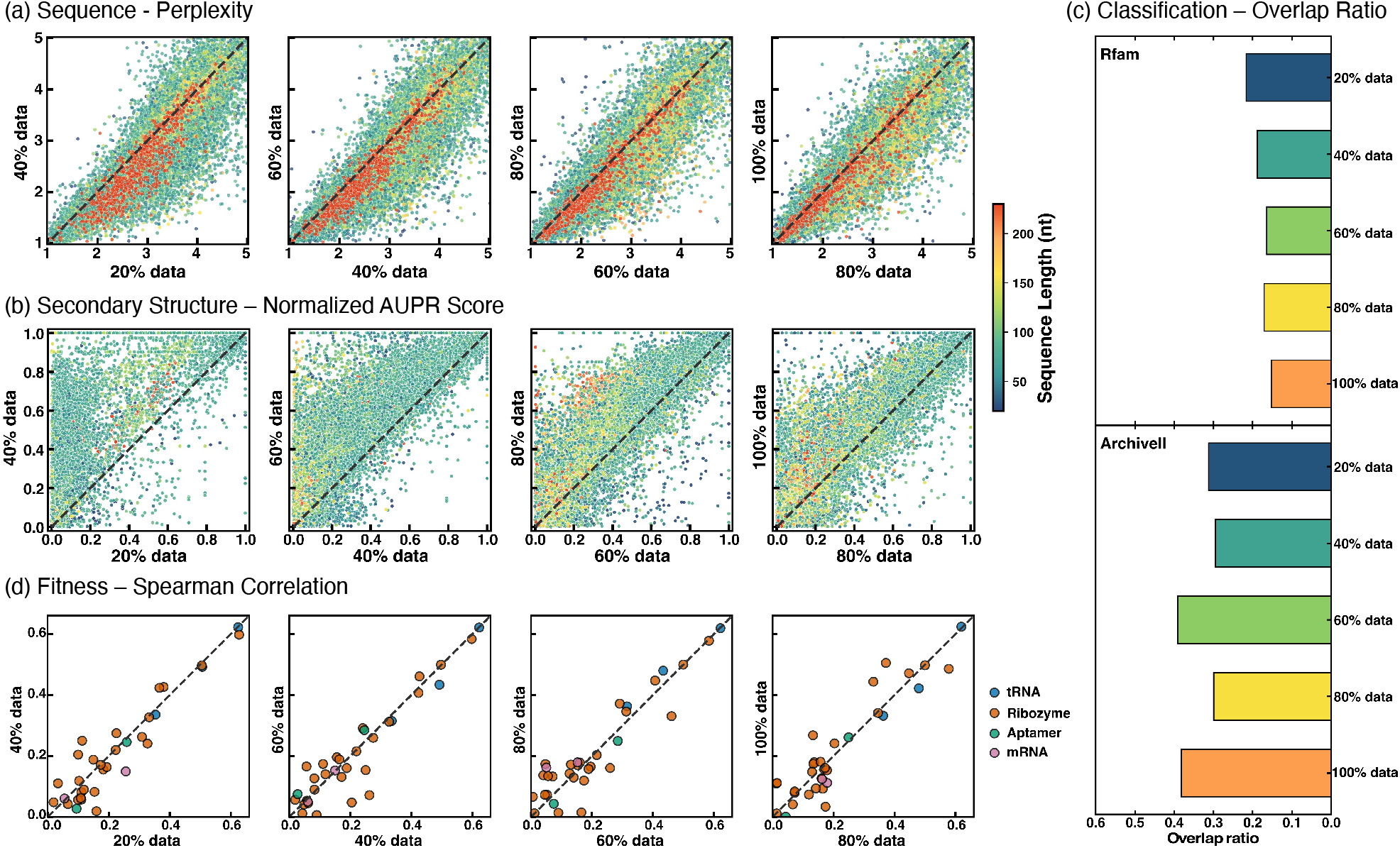
Emergence of RNA structural and functional representations with training-data scaling. The 100M-parameter LIMBER-*nano* model is trained for six epochs using progressively larger fractions of the training dataset. Performance is evaluated using the same benchmarks and metrics described in Extended Fig. 1. (a) Sequence-level perplexity on bpRNA-1m(90). (b) Zero-shot secondary-structure prediction performance. (c) Zero-shot RNA classification for homologous and non-homologous sequences sampled from the Rfam and ArchiveII datasets. (d) Zero-shot RNA fitness prediction across 33 deep mutational scanning assays.

