## Supplemental Information for "LIMBER: A Hybrid RNA Foundation Model for Predicting and Understanding RNA Structure and Function"

**Contents**

I. Overview of LIMBER Model 1
II. Model Architectures and Sequence Dataset for Pre-training 2
III. Zero-shot Secondary Structure Prediction: Categorical Jacobian and Embedding Perturbation 4
IV. Fine-tuning LIMBER for Secondary Structure Prediction 5
V. Fine-tuning LIMBER for ncRNA Classification 6
VI. Internal Ribosome Entry Site Identification 7
VII. Fine-tuning LIMBER for Mean Ribosome Load Prediction 8
VII. LIMBER-Driven Prediction of RNA–Small-Molecule Interactions 8

**References** 11

### I. Overview of LIMBER Model

LIMBER (Layer-Interleaved Mamba-Bidirectional Encoder for RNA) is a hybrid RNA foundation model that integrates bidirectional self-attention[1] with state-space sequence modeling[2, 3] within a unified masked-language-modeling encoder. Given an RNA sequence tokenized at single-nucleotide resolution, LIMBER first maps the input sequence into continuous representations using learned nucleotide embeddings and absolute positional embeddings. The resulting hidden representations are then processed by a stack of Transformer and Mamba layers interleaved according to a repeating Transformer-Transformer-Mamba (TTM) pattern, such that approximately one-third of the encoder layers are Mamba blocks and the remaining layers are bidirectional Transformer blocks.

Each Transformer block consists of multi-head self-attention followed by a position-wise feed-forward network, with residual connections, layer normalization, GELU activations, and dropout. Self-attention is bidirectional, allowing each nucleotide representation to directly integrate contextual information from the complete input sequence. Differently, each Mamba block applies a selective state-space operator within a pre-layer-normalized residual formulation,

$$\mathbf{H}^{(l+1)} = \mathbf{H}^{(l)} + \text{Dropout}\left[\text{Mamba}\left(\text{LN}\left(\mathbf{H}^{(l)}\right)\right)\right], \tag{1}$$

where $\mathbf{H}^{(l)}$ denotes the hidden representation entering layer $l$. All Mamba layers use a state dimension $d_{\text{state}} = 16$, convolution width $d_{\text{conv}} = 4$, and expansion factor of 4. Individual Mamba layers operate causally from left to right, whereas the interleaved Transformer layers provide global bidirectional context mixing throughout the encoder. Input sequences are right-padded; consequently, trailing padding tokens do not affect representations of preceding valid nucleotides during the causal Mamba scan.

---

The final encoder representations are passed to a masked-language-modeling prediction head consisting of a linear transformation, GELU activation, layer normalization, and a final linear projection to the nucleotide vocabulary. The output decoder weights are not tied to the input embedding matrix. LIMBER uses a 10-token vocabulary comprising five special tokens (`[PAD]`, `[UNK]`, `[CLS]`, `[SEP]`, and `[MASK]`), the four canonical RNA nucleotides (`A`, `U`, `C`, and `G`), and an ambiguous nucleotide token `N`. During sequence preprocessing, all nucleotides are converted to uppercase, thymine is replaced with uracil, and non-`ACGU` characters are mapped to `N`.

LIMBER is pretrained using masked language modeling. For each input sequence, 15% of nucleotide positions are selected for prediction. Among the selected positions, 80% are replaced by the `[MASK]` token, 10% are replaced by a randomly sampled canonical nucleotide from {`A`, `U`, `C`, `G`}, and the remaining 10% are left unchanged. Cross-entropy loss is computed only over the selected positions. Through this objective, LIMBER learns contextual representations directly from unlabeled RNA sequences while its interleaved architecture combines bidirectional attention-based context integration with state-space sequence modeling.

To systematically investigate model scaling and assess the advantages of the hybrid architecture, we train LIMBER variants spanning five model sizes (10M, 50M, 100M, 300M and 650M parameters), four additional LIMBER-100M variants using 20%, 40%, 60% and 80% of the full pretraining dataset, and a LIMBER-100M variant with a reduced causal convolution kernel width of 2, yielding a total of 10 LIMBER variants. All variants retain the same TTM interleaving architecture. For architectural comparisons, we additionally train pure BERT and Mamba models at three matched parameter scales (100M, 300M and 650M). Detailed architectural configurations, parameterization, pretraining data, and optimization settings are provided in Sec. II.

### II. Model Architectures and Sequence Dataset for Pre-training

***Model Configurations.*** We pretrain 16 models encompassing the hybrid LIMBER architecture, its pure BERT and Mamba counterparts, and model- and data-scaling variants: 10 LIMBER variants, three BERT models and three Mamba models. Their configurations are summarized in Table S1, and all 16 pretrained models are publicly available. Every model shares the same nucleotide vocabulary, masking objective, sequence preprocessing procedure and maximum context length, so that architecture, model capacity and pretraining-data volume are the only quantities that vary. The maximum input length is 1,022 nucleotides. In the main text, LIMBER-100M, LIMBER-300M and LIMBER-650M are referred to as LIMBER-*nano*, LIMBER-*meso* and LIMBER-*macro*, respectively.

Within the LIMBER series, model capacity is scaled by increasing the hidden dimension $d_{\mathrm{model}}$, which determines the width of interlayer representations, and by increasing encoder depth at a fixed hidden dimension. The parameter count of the attention and feed-forward components of a Transformer layer grows approximately quadratically with $d_{\mathrm{model}}$, so a layer of width 1,280 holds roughly four times as many parameters as a layer of width 640. LIMBER-*nano* comprises 91.7M parameters across 18 encoder layers with a hidden dimension of 640, whereas LIMBER-*meso* contains 303.6M parameters across 15 layers with a doubled hidden dimension of 1,280. The reduced encoder depth partially offsets the increase in parameter count associated with the wider representations. LIMBER-*meso* and LIMBER-*macro* share the same hidden dimension, number of attention heads and feed-forward dimension, so their difference in size is determined by encoder depth alone. These variants enable systematic analyses along three dimensions. The series spanning LIMBER-10M, LIMBER-50M, LIMBER-*nano*, LIMBER-*meso* and LIMBER-*macro* examines scaling with model capacity; four LIMBER-*nano* variants pretrained on 20%, 40%, 60% and 80% of the full corpus assess the effects of pretraining data volume; and LIMBER-*nano-Conv2*, which differs from LIMBER-*nano* only in the reduced Mamba causal convolution kernel width (from 4 to 2), probes the contribution of local sequential context.

Every third encoder layer of a LIMBER model is a Mamba layer. All Mamba layers use a state dimension of $d_{\mathrm{state}} = 16$, a convolution width of $d_{\mathrm{conv}} = 4$ and an expansion factor of 4; the single exception is LIMBER-*nano-Conv2*, which reduces the convolution width to $d_{\mathrm{conv}} = 2$. Dropout is set to 0.1 in all encoder layers of all architectures.

The BERT and Mamba controls match the corresponding LIMBER models in encoder depth and hidden dimension, and the BERT controls additionally match the number of attention heads and the feed-forward dimension. Every encoder layer is a Transformer layer in the BERT models and a Mamba layer in the Mamba models, so that a comparison at a fixed parameter scale attributes differences in performance to the composition of layer types rather than to model size or training budget. Algorithm 1 describes the construction of an encoder from a specified layer

Table S1: **Configurations of the sixteen pretrained models.** $L$ denotes the number of encoder layers; $d_{\text{model}}$, the hidden dimension; Heads, the number of self-attention heads; and $d_{\text{ff}}$, the feed-forward dimension of Transformer layers. In the layer pattern, T and M denote bidirectional Transformer and Mamba layers, respectively, with (TTM) $\times$ $n$ indicating $n$ repetitions of the interleaved TTM block. Attention heads and Transformer feed-forward dimensions are not applicable to Mamba layers and are therefore marked with a dash. Data indicates the fraction of the training corpus (40,498,819 deduplicated sequences, excluding the held-out validation split) on which each model was pretrained. Parameter counts include token and positional embeddings and the masked-language-modeling head. Unless otherwise specified, all Mamba layers use a state dimension of $d_{\text{state}} = 16$, a causal convolution kernel width of $d_{\text{conv}} = 4$ and an expansion factor of 4.

| Model | Layer pattern | $L$ | $d_{\text{model}}$ | Heads | $d_{\text{ff}}$ | Data | Parameters |
|---|---|---|---|---|---|---|---|
| *LIMBER (interleaved Transformer–Transformer–Mamba)* | | | | | | | |
| LIMBER 10M | (TTM) × 4 | 12 | 256 | 4 | 1,024 | 100% | 10.2M |
| LIMBER 50M | (TTM) × 5 | 15 | 512 | 8 | 2,048 | 100% | 49.3M |
| LIMBER-*nano* | (TTM) × 6 | 18 | 640 | 10 | 2,560 | 100% | 91.7M |
| LIMBER-*meso* | (TTM) × 5 | 15 | 1,280 | 20 | 5,120 | 100% | 303.6M |
| LIMBER-*macro* | (TTM) × 11 | 33 | 1,280 | 20 | 5,120 | 100% | 664.3M |
| LIMBER-*nano*-20% | (TTM) × 6 | 18 | 640 | 10 | 2,560 | 20% | 91.7M |
| LIMBER-*nano*-40% | (TTM) × 6 | 18 | 640 | 10 | 2,560 | 40% | 91.7M |
| LIMBER-*nano*-60% | (TTM) × 6 | 18 | 640 | 10 | 2,560 | 60% | 91.7M |
| LIMBER-*nano*-80% | (TTM) × 6 | 18 | 640 | 10 | 2,560 | 80% | 91.7M |
| LIMBER-*nano-Conv2*† | (TTM) × 6 | 18 | 640 | 10 | 2,560 | 100% | 91.7M |
| *BERT (all Transformer layers)* | | | | | | | |
| BERT 100M | T × 18 | 18 | 640 | 10 | 2,560 | 100% | 89.7M |
| BERT 300M | T × 15 | 15 | 1,280 | 20 | 5,120 | 100% | 298.1M |
| BERT 650M | T × 33 | 33 | 1,280 | 20 | 5,120 | 100% | 652.3M |
| *Mamba (all Mamba layers)* | | | | | | | |
| Mamba 100M | M × 18 | 18 | 640 | – | – | 100% | 95.8M |
| Mamba 300M | M × 15 | 15 | 1,280 | – | – | 100% | 314.4M |
| Mamba 650M | M × 33 | 33 | 1,280 | – | – | 100% | 688.2M |

† Causal convolution kernel width reduced to $d_{\text{conv}} = 2$; all other settings match LIMBER-*nano*.

pattern and the propagation of input sequences through the resulting architecture. The three architectures differ only in the layer pattern $P$.

***Dataset Construction.*** The pretraining dataset is derived from RNAcentral[4], released on 14 May 2025. We apply no filter on RNA family, sequence length or sequence identity, retaining the full active set rather than the homology-reduced subsets used by several earlier RNA language models. The only reduction is exact duplicate removal, in which sequences that are identical after case folding are collapsed to a single representative, leaving 40,518,474 unique sequences. Each sequence is then mapped onto the four-letter RNA alphabet by converting thymine to uracil and every remaining non-`ACGU` character, including all IUPAC ambiguity codes, to `N`. A fixed validation set of 19,655 sequences is selected from the deduplicated collection, leaving 40,498,819 sequences for pretraining, and the same validation set is used throughout. Each training epoch therefore contains all 40,498,819 deduplicated sequences, corresponding to approximately 18.85 billion nucleotide tokens. Sequences longer than 1,022 nucleotides are represented by a randomly selected contiguous subsequence in each epoch: for a sequence of length $L > 1{,}022$, the starting position $s$ is sampled uniformly from the valid starting positions, $s \sim \mathcal{U}(0, L - 1{,}022)$, and the 1,022-nucleotide subsequence beginning at position $s$ is used as the model input. Because a new starting position is sampled in each epoch, different regions of a long RNA are observed over successive epochs. Sequences of at most 1,022 nucleotides are retained in full and right-padded within each batch. The fixed validation set is used for all different models and training epochs. The same pretraining corpus and the same held-out validation set are shared across all model variants. Models trained on the full corpus see an identical training set, and the four data-scaling variants are drawn as nested, category-stratified subsets of that same corpus, such that the 20% subset is contained in the 40% subset, which is contained in the 60% subset, and so on. Validation sequences are excluded from the training data of every variant.

***Pretraining Optimization.*** All models, including the LIMBER, BERT and Mamba variants used in the scaling analysis and benchmarks, are pretrained with identical optimization settings. We adopt the AdamW optimizer

**Algorithm 1** Encoder construction and forward pass under a layer pattern

**Input:** Layer pattern $P \in \{\mathrm{T}, \mathrm{M}\}^L$; token indices $x \in \{1, \ldots, V\}^N$; padding mask $m \in \{0, 1\}^N$

**Construction**

**for** $\ell = 1$ **to** $L$ **do**

$$\text{layer}_\ell \leftarrow \begin{cases} \textsc{TransformerBlock}(d_{\text{model}}, \text{Heads}, d_{\text{ff}}) & P_\ell = \mathrm{T} \\ \textsc{MambaBlock}(d_{\text{model}}, d_{\text{state}}, d_{\text{conv}}, \text{expand}) & P_\ell = \mathrm{M} \end{cases}$$

**end for**

**Forward pass**

$H \leftarrow \textsc{TokenEmbed}(x) + \textsc{PositionEmbed}(1{:}N)$

$H \leftarrow \textsc{Dropout}(\textsc{LayerNorm}(H))$

$M_{\text{attn}} \leftarrow (1 - m) \cdot (-\infty)$ ▷ additive mask; padded positions are excluded from attention

**for** $\ell = 1$ **to** $L$ **do**

  **if** $P_\ell = \mathrm{T}$ **then**

    $H \leftarrow \textsc{LayerNorm}\big(H + \textsc{SelfAttention}(H, M_{\text{attn}})\big)$

    $H \leftarrow \textsc{LayerNorm}\big(H + \textsc{FeedForward}(H)\big)$ ▷ bidirectional, padding-aware

  **else**

    $H \leftarrow H + \textsc{Dropout}\big(\textsc{SelectiveSSM}(\textsc{LayerNorm}(H))\big)$ ▷ pre-norm residual; causal along $5' \rightarrow 3'$

  **end if**

**end for**

**return** $\textsc{MLMHead}(H) \in \mathbb{R}^{N \times V}$

($\beta_1 = 0.9$, $\beta_2 = 0.999$, $\epsilon = 10^{-8}$) with a weight decay of 0.01. The learning rate is warmed up linearly over the first 2,000 steps to a peak of $5 \times 10^{-5}$ followed by a cosine schedule decaying to a minimum of $1 \times 10^{-5}$. Gradients are clipped to a maximum norm of 1.0, training uses bfloat16 mixed precision, and the random seed is fixed to 42. Dropout is set to 0.1 in all encoder layers for all architectures. The effective batch size is 2,688 sequences for every model, obtained by combining the per-device batch size with gradient accumulation across devices. Pretraining is performed on four NVIDIA H200 GPUs, and all fine-tuning and downstream experiments are performed on four NVIDIA L40S GPUs.

### III. Zero-shot Secondary Structure Prediction: Categorical Jacobian and Embedding Perturbation

***Dataset Construction.*** We adopt the integrative bpRNA-1m(90) dataset[5], which consolidates RNA sequences and structures from multiple sources, including the Comparative RNA Web (CRW), tmRNA Database, tRNAdb 2009, Signal Recognition Particle (SRP) Database, RNase P Database, the RCSB Protein Data Bank (PDB), and all RNA families in Rfam 12.2. During dataset construction, the sequences with > 90% sequence similarity when there is at least 70% alignment coverage are removed, and sequences longer than 512 nucleotides are removed, resulting in 26,707 sequences (see Tabel S2). In the categorical Jacobian and embedding perturbation inference, the pseudoknots in the labeled secondary structures are regarded as normal based pairs.

Table S2: Number of sequences in refined bpRNA_1m(90) datasets[5]

| Dataset | CRW | PDB | RFAM | RNP | SRP | tRNAdb | tmRNA |
|---|---|---|---|---|---|---|---|
| # of Sequences | 3,300 | 329 | 21,931 | 253 | 348 | 207 | 339 |

The following provides a brief description for each dataset, including the approaches used to generate secondary structure annotations:
(1). CRW secondary structure annotations are derived from comparative sequence analysis of homologous RNAs, in which conserved base-pairing interactions are inferred from patterns of nucleotide covariation, including compensatory substitutions. These interactions are integrated into comparative structural models and transferred to individual RNA sequences through sequence-structure alignments, with additional refinement where necessary.
(2). The tm-RNA Database includes the sequences and structures of transfer messenger RNAs (tmRNAs), which are bacterial RNAs with both tRNA- and mRNA-like functions. Their secondary structures are also inferred through

comparative sequence analysis. The structure annotations are primarily derived from computational analyses, with partial experimental validation.
(3). Signal Recognition Particle (SRP) Database is a curated database of sequences associated with the structure and function of signal recognition particle RNAs, where the secondary structures are primarily derived through comparative sequence analysis.
(4). tRNAdb 2009 dataset, formerly known as the Sprinzl tRNA Database, is jointly maintained by research groups in Leipzig, Marburg, and Strasbourg. It contains 623 experimentally characterized tRNA sequences from 104 organisms. The corresponding secondary structures are curated cloverleaf annotations derived from established tRNA structural conventions, standardized sequence alignments, prior experimental evidence, and gene annotations.
(5). The RNase P Database (RNP) primarily covers bacterial and archaeal RNase P RNAs, and the corresponding secondary structures are curated, phylogenetically informed models derived from homologous-sequence alignments and established RNase P structural frameworks.
(6). RCSB Protein Data Bank (PDB) subset includes all entries deposited before June 12, 2017, that contain at least one RNA chain. RNAView was used to identify base-pairing interactions from the experimentally determined three-dimensional coordinates, and the resulting secondary structures are converted into BPSEQ format.
(7). The RNA Family Database (RFAM) V12.2 contains consensus structures derived from comparative sequence analysis of individual sequence family members of thousands of RNA families.

Some prior studies have used the ArchiveII dataset (3,865 sequences) to benchmark RNA secondary-structure prediction. Despite having distinct record identifiers, ArchiveII contains duplicate sequences. The refined bpRNA-1m(90) dataset contains no redundant sequences. We note that approximately 41% of ArchiveII sequences overlap with the refined bpRNA-1m(90) dataset, with particularly high sequence similarity ($\geq 97\%$) observed for the RNP, SRP and tRNAdb subsets, alongside substantial overlap in the tmRNA subset.

### IV. Fine-tuning LIMBER for Secondary Structure Prediction

We evaluate LIMBER on supervised RNA secondary structure prediction by fine-tuning the pretrained model to predict nucleotide pairing matrices. Given an RNA sequence of length $L$, the model predicts an $L \times L$ matrix in which each entry represents the pairing score between two nucleotide positions. This supervised setting complements the task-agnostic structural analyses described in the main text by allowing the pretrained representations to be adapted directly using annotated RNA secondary structures.

***Dataset Construction.*** Supervised secondary-structure prediction is evaluated on three benchmarks that probe complementary notions of generalization: an intra-family benchmark derived from bpRNA, a benchmark whose test structures are deliberately distinct from those seen in training, and a cross-family leave-one-family-out benchmark. Across all three, sequences longer than 512 nucleotides are excluded, and every model is trained and evaluated on identical splits. Detailed descriptions of the datasets are provided below.

*bpRNA (TR0/VL0/TS0)*[5]. We use the curated bpRNA splits provided by Singh et al.[6], comprising 13,419 RNA sequences partitioned into training (TR0; 10,814 sequences), validation (VL0; 1,300 sequences) and independent test (TS0; 1,305 sequences) sets, with sequence redundancy reduced at an 80% identity threshold. All sequences are within the 512-nt length limit and are therefore retained for evaluation.

*Rivas (TrainSetA, TestSetA and TestSetB).* We additionally adopt the benchmark from [7], distributed with MXfold2 [8]. TrainSetA aggregates 3,166 sequences from published RNA structure benchmarks; 223 exceed 512 nucleotides and are removed, and the remaining 2,943 are split at random into 2,649 training and 294 validation sequences. TestSetA contains 547 sequences after the same length restriction removes 45 of its 592 sequences. TestSetB contains 430 sequences drawn from Rfam v10.0 seed alignments [9], none of which exceed 512 nucleotides, so the length restriction removes nothing and the full set is retained. Their structures are distinct from those represented in TrainSetA, thereby assessing generalization to structurally divergent RNAs.

*ArchiveII (leave-one-family-out).* Cross-family transferability is evaluated on the ArchiveII dataset[10] following the protocol of [11] across nine RNA families: 5S rRNA, 16S rRNA, 23S rRNA, Group I intron, SRP RNA, Telomerase RNA, RNaseP RNA, tmRNA and tRNA. Sequences are restricted to 30-512 nucleotides, exact duplicates are removed, and each family is capped at 200 randomly sampled sequences to reduce class imbalance, as family sizes in the original dataset vary by more than two orders of magnitude. This yields 1,188 RNAs in total: 200 each for 5S rRNA, SRP RNA, RNase P RNA, tmRNA and tRNA, and 71, 67, 35 and 15 for Group I intron, 16S rRNA, Telomerase RNA and 23S rRNA, respectively. For each of the nine evaluation folds, one family is held out as the test set and the remaining eight provide the training and validation data, so that each family serves exactly once as the independent test family.

***Secondary Structure Prediction Head.*** The secondary structure prediction head follows the architecture used in RiNALMo[12]. Let $\mathbf{H} = (\mathbf{z}_1, \mathbf{z}_2, \ldots, \mathbf{z}_L), \mathbf{z}_i \in \mathbb{R}^d$; denote the final layer representations produced by the pretrained encoder. Pairwise features are constructed by concatenating the representations at nucleotide positions $i$ and $j$, as $\mathbf{p}_{ij} = [\mathbf{z}_i; \mathbf{z}_j] \in \mathbb{R}^{2d}$. Applying this operation to all nucleotide pairs produces a tensor of dimension $L \times L \times 2d$ for each sequence. Each pairwise representation is first projected from $2d$ dimensions to 64 channels using a linear transformation. The resulting pairwise feature map is processed by two residual 2D convolutional blocks. Each block consists of a $1 \times 1$ convolution followed by instance normalization and ReLU activation, a $3 \times 3$ convolution with same padding followed by instance normalization and ReLU activation, and a final $1 \times 1$ convolution followed by instance normalization and ReLU activation. The output of the convolutional sequence is added to the input of the block through a residual connection. A final $3 \times 3$ convolution maps the 64-channel representation to a single pairing logit for each pair of nucleotide positions. The resulting matrix is symmetrized to obtain a single score for each unordered nucleotide pair. Only entries above the main diagonal are used during training and evaluation.

***Fine-Tuning Protocol.*** We follow the gradual unfreezing procedure used for RiNALMo[12]. At the beginning of fine-tuning, all pretrained encoder parameters are frozen and only the structure prediction head is optimized. Every three epochs, the next three encoder layers are unfrozen in order from the top of the encoder toward the input. Newly unfrozen encoder layers are added to the optimizer with a learning rate ten times smaller than that of the prediction head.

The lowest encoder layers remain frozen throughout fine-tuning. For the 100M and 300M models, the bottom three layers are permanently frozen, and for the 650M models, the bottom nine layers are permanently frozen. Fine-tuning is performed for 30 epochs. The Adam optimizer is consistently employed. The learning rates for the prediction head and unfrozen encoder layers are set to $5 \times 10^{-4}$ and $5 \times 10^{-5}$, respectively. The learning rate is linearly decayed from a multiplicative factor of 1.0 to 0.1 over 7,000 optimization steps. The training objective is binary cross entropy computed over entries in the upper triangular portion of the predicted pairing matrix. For a predicted logit matrix $\mathbf{S}$ and binary reference contact matrix $\mathbf{Y}$, the loss is evaluated as:

$$\mathcal{L}_{\mathrm{BCE}} = -\frac{1}{|\Omega|} \sum_{(i,j)\in\Omega} \left[ Y_{ij} \log \sigma(S_{ij}) + (1 - Y_{ij}) \log \left(1 - \sigma(S_{ij})\right) \right], \tag{2}$$

where $\Omega = \{(i, j) : j > i\}$ and $\sigma(\cdot)$ denotes the sigmoid function. No positive class weighting is applied.

***Inference and Structure Decoding.*** The decision threshold for base pair prediction is selected using the validation set. We evaluate thresholds from 0.01 to 0.29 in increments of 0.01 and select the value that maximizes validation F1. Predicted base pairs are decoded using greedy maximum probability matching. Candidate nucleotide pairs are ranked by their predicted probabilities and processed in descending order. A candidate pair is retained only if neither nucleotide has previously been assigned to another pair. We further restrict predictions to canonical and wobble base pairs, $\{\mathrm{AU}, \mathrm{UA}, \mathrm{GC}, \mathrm{CG}, \mathrm{GU}, \mathrm{UG}\}$, and exclude sharp loops satisfying $|i - j| < 4$.

***Evaluation.*** Secondary structure prediction is evaluated using the flexible F1 score, in which a predicted base pair is considered correct when it matches a reference pair within a tolerance of one nucleotide position. The score is computed per sequence. Four published RNA language models serve as baselines, spanning the two parameter scales of interest: RiNALMo-giga and AIDO.RNA-650M at approximately 650M parameters, and RNA-FM and structRFM at approximately 100M parameters. Every baseline is fine-tuned and evaluated with the same prediction head, gradual unfreezing schedule, decoding procedure and evaluation protocol as LIMBER, and the number of permanently frozen bottom layers follows encoder depth rather than model name: nine for the 33-layer encoders (LIMBER-*macro*, RiNALMo-giga and AIDO.RNA-650M) and three for the shallower ones (LIMBER-*nano*, LIMBER-*meso*, RNA-FM and structRFM). The pretrained backbone is therefore the only quantity that differs between runs, and differences in performance reflect differences in the learned RNA representations.

### V. Fine-tuning LIMBER for ncRNA Classification

***Dataset Construction.*** We employ the nRC benchmark[13], which assigns ncRNAs to 13 functional classes: 5S rRNA, 5.8S rRNA, CD-box and HACA-box snoRNAs, group I and group II introns, IRES, leader sequences, miRNA, ribozymes, riboswitches, scaRNA and tRNA. The training and evaluation sets comprise 6,320 and 2,600 sequences, respectively, with the latter balanced at 200 sequences per class. Sequences range from 38 to 1,136 nucleotides with a median of 130; inputs are truncated at 1,022 nucleotides, which affects a single training sequence.

***Fine-Tuning and Prediction.*** Each pretrained language model serves as the sequence encoder, with all downstream components held fixed to ensure that the encoder is the sole variable in the comparison. The `[CLS]` representation from the final encoder layer is passed through layer normalization and dropout 0.1 to a single linear projection onto the 13 classes. Fine-tuning proceeds in two phases: the encoder is initially frozen while the prediction head is optimized for 10 epochs using AdamW (learning rate $5 \times 10^{-4}$, weight decay 0.01), followed by end-to-end fine-tuning for an additional 20 epochs with reduced learning rates for the encoder (encoder learning rate $1 \times 10^{-5}$, head $1 \times 10^{-4}$). All runs use cross-entropy loss, an effective batch size of 64 sequences achieved through gradient accumulation, gradient clipping at a maximum norm of 1.0, and a fixed random seed of 42. Model weights are updated on the training set only; the checkpoint with the highest validation accuracy is retained, and the same protocol is applied to every model. RiNALMo Giga, AIDO.RNA-650M, RNA-FM and StructRFM are fine-tuned through the same head, optimization and evaluation pipeline.

***Encoder-Depth Ablation.*** Our layer-wise analyses reveal that discriminative information about ncRNA families and types is strongly accumulated in intermediate encoder layers, with LIMBER-macro exhibiting a low distribution overlap ratio at layer 14. Motivated by this observation, we perform an encoder-depth ablation in which LIMBER-macro is truncated to the first 14 of its 33 encoder blocks, corresponding to the interleaving pattern `TTMTTMTTMTTMTT`, and, as a lower reference point, to the first 2 blocks, corresponding to `TT`. Each truncated model uses the final retained layer's `[CLS]` representation for classification and is fine-tuned with the protocol applied to the full model.

***Evaluation.*** Class predictions are assigned based on the highest predicted probability. We report accuracy, Matthews correlation coefficient (MCC), F1 score, AUROC and AUPRC, with the latter two computed using a one-versus-rest strategy across all 13 classes. Corresponding class-specific F1 scores are also reported. Accuracy, MCC and F1 follow from the arg-max over the 13 class probabilities.

### VI. Internal Ribosome Entry Site Identification

***Dataset Construction.*** We employ the DeepIRES benchmark [14] and its released split files without modification. The dataset is constructed via a high-throughput bicistronic reporter assay of ~55,000 designed oligonucleotides spanning known IRESs, human and viral genomic fragments, and synthetic sequences [15]. Following [14], the screen is filtered in three steps: synthetic sequences are discarded and only native sequences from eight screen categories retained (Genome_Wide_Screen_Elements, IRESite_blocks, Viral_5UTR_Screen and five others); sequences with promoter activity $> 0.2$ or splicing activity $< -2.5$ in the companion assays of the same experiment are removed as artifacts; and CD-HIT-EST is applied at an identity cutoff of 0.8. The resulting dataset comprises 18,527 sequences, each 174 nt in length, with 1,917 sequences exhibiting measured IRES activity $> 600$ labeled as positive and the remaining 16,610 as negative. The dataset is partitioned into 16,674 training sequences (90%) and 1,853 held-out test sequences (10%), containing 1,728 (10.4%) and 189 (10.2%) positive sequences, respectively.

For additional evaluation, we use the main independent test set compiled by DeepIRES [14], comprising 104 sequences with equal numbers of positive and negative examples. Specifically, 40 positive and 40 negative sequences are drawn from the IRESfinder independent test set [16], while an additional 12 positive and 12 negative sequences are obtained from the experimental study of Wellensiek et al. [17]. Sequence lengths range from 54 to 174 nt, compared with the fixed length of 174 nt in the DeepIRES training set. Exact sequence matching identifies 34 sequences shared with the DeepIRES training partition.

***Fine-Tuning and Prediction.*** Each pretrained RNA language model serves as the sequence encoder, with the prediction head, datasets and optimization settings held constant to ensure that the encoder is the sole variable in the comparison. Hidden states from the final encoder layer are mean-pooled over nucleotide positions, excluding `[CLS]`, `[SEP]` and padding. The pooled representation of dimension $d$ is processed through layer normalization, dropout (rate 0.1), and a linear projection to $d/2$ dimensions, followed by GELU activation and a final linear layer that produces a single classification logit.

A random 10% of the training partition (seed 42) is held out for model selection, leaving 15,007 training and 1,667 validation sequences. Training proceeds in two phases. The encoder is first frozen and the head alone is trained for 8 epochs with AdamW (learning rate $5 \times 10^{-4}$, weight decay 0.01); the head state with the highest validation AUROC is then restored, the encoder is unfrozen, and the full model is trained for a further 12 epochs with AdamW (encoder learning rate $1 \times 10^{-5}$, head $1 \times 10^{-4}$, weight decay 0.01). Both phases use batch size 32 and gradients clipped to a maximum norm of 1.0. The objective is binary cross-entropy on the logit, weighted by the negative-to-positive ratio of the training split to offset the 1:9 class imbalance. The full-model state with the highest validation AUROC is retained and used for every reported test number. All models share random seed 42. RiNALMo-giga, AIDO.RNA-650M, RNA-FM and structRFM are fine-tuned and evaluated using identical pooling strategies, prediction heads, optimization protocols and evaluation metrics. The task-specific methods DeepIRES [14] and IRESfinder [16] are included as additional benchmarks on the same test sets.

***Evaluation.*** Performance is evaluated using two threshold-independent metrics, AUROC and AUPRC, and three threshold-dependent metrics–accuracy, F1 score and Matthews correlation coefficient (MCC)–calculated at a fixed decision threshold of 0.5. All metrics are computed consistently across the two test sets.

### VII. Fine-tuning LIMBER for Mean Ribosome Load Prediction

***Dataset Construction.*** We predict mean ribosome load (MRL) from 5′UTR sequence using the variable-length massively parallel reporter assay of Sample et al.[18], in the Random_Vary and Human_Vary split of the Karollus benchmark. Models are trained on the 76,319 random-library UTRs and evaluated on two held-out sets of 7,600 sequences each, one drawn from the same random library and one from human 5′UTRs. All sequences are 25–100 nucleotides, with the two test sets containing exactly 100 sequences at each length. MRL values span a mean of 5.27 (s.d. 1.36) in training and 5.20 (s.d. 1.41) and 5.54 (s.d. 1.20) in the random and human test sets.

***Fine-Tuning and Prediction.*** Each pretrained language model is used as the sequence encoder with everything downstream held fixed. Final-layer hidden states are mean-pooled over nucleotide positions, excluding special and padding tokens, and passed to a regression head consisting of layer normalization, dropout 0.1, a linear projection to half the hidden width, a GELU nonlinearity and a linear projection to a scalar. Fine-tuning proceeds in two phases: the encoder is frozen and the head trained alone for 8 epochs with AdamW (learning rate $5 \times 10^{-4}$, weight decay 0.01), after which the encoder is unfrozen and the full model trained for a further 12 epochs (encoder learning rate $1 \times 10^{-5}$, head $1 \times 10^{-4}$). Both training phases use a batch size of 64, Huber loss with $\delta = 1.0$, gradient clipping at a maximum norm of 1.0, and a fixed random seed of 42. A random 10% of the training set is held out for checkpoint selection on validation $R^2$, leaving 68,688 training and 7,631 validation sequences. RiNALMo Giga, AIDO.RNA-650M, RNA-FM and StructRFM are fine-tuned through the identical pooling, head, optimization and evaluation pipeline, differing only in the encoder.

***Evaluation.*** Performance is quantified by the squared Pearson correlation coefficient $R^2$ between predicted and experimentally measured MRL, evaluated separately on the random and human test sets. All models, including the baselines, are scored with the same function, so the comparison between them is unaffected by this choice.

### VII. LIMBER-Driven Prediction of RNA–Small-Molecule Interactions

***Dataset Construction.*** We use the SMRTnet dataset reported in [19] without modification. It is derived from 1,061 RNA–small-molecule complex structures in the Protein Data Bank (PDB): experimentally resolved interactions provide the positive pairs, and negative pairs are generated by randomly pairing RNAs and small molecules after excluding known interactions, giving 24,687 pairs at a 1:2 positive-to-negative ratio. Following the original protocol, each RNA is represented by the 31-nucleotide fragment centered on its binding pocket together with the corresponding experimentally derived secondary structure, and each small molecule by its SMILES string and molecular graph.

Training, validation and test partitions are the five-fold cross-validation splits released with SMRTnet, read from fixed files rather than resampled: the folds are defined by SMILES, so no small molecule in a test fold appears in the corresponding training or validation fold. All RNA encoders and random seeds are evaluated on the same five predefined folds, ensuring consistent data partitions across comparisons, with random seeds affecting only training stochasticity. Transferability is assessed on the SMRTnet benchmark dataset, assembled from four resources (R-BIND, R-SIM, SMMRNA and NALDB) and 22 publications, from which interactions overlapping the SMRTnet dataset have been removed. It comprises 2,011 experimentally characterized RNA–small-molecule interactions, 1,665 positive and 346 negative.

***Model Configuration.*** We adopt the SMRTnet architecture and replace only its RNA language model, RNASwan-seq, with each RNA language model under comparison. The chemical language model MoLFormer [20], the convolutional secondary-structure encoder, the graph-attention encoder for molecular structure, the co-attention multimodal fusion module and the classification head are the original SMRTnet modules and are left untouched, so that the RNA encoder is the only quantity that differs between runs. All modules other than the two pretrained language models are randomly initialized.

The substituted encoder must satisfy the interface SMRTnet expects of RNASwan-seq: given `[CLS]` followed by 31 nucleotide tokens, it returns a $32 \times 640$ representation from which the `[CLS]` position is dropped, leaving one 640-dimensional vector per nucleotide. Encoders whose hidden width is already 640 (LIMBER-*nano* and RNA-FM) feed the downstream stack directly, whereas encoders of width 1,280 (LIMBER-*macro*, RiNALMo-giga and AIDO.RNA-650M) pass through a single learned linear projection to 640 dimensions. The projection is the only parameter added to the published architecture, and it ensures that every backbone feeds an identical head. Each backbone uses its own nucleotide vocabulary, so token indices are remapped explicitly rather than by retokenizing. Ambiguity codes, which SMRTnet encodes but our single-nucleotide vocabulary does not represent, are mapped to `N`. Two backbone-specific conventions are handled in the same place: our models are pretrained with a trailing `[SEP]` token that SMRTnet's tokenizer never emits, which is therefore appended before the forward pass and stripped from the output; RiNALMo uses a DNA alphabet containing no `U`, so uracil is mapped to thymine.

The remaining architectural hyperparameters follow those of the original published model. Specifically, the secondary-structure encoder employs 16 channels with a kernel size of 7 and padding of 3, while the molecular graph-attention encoder comprises three layers with 74 input features, a hidden dimension of 256, three attention heads, and a dropout rate of 0.2 applied to both node features and attention coefficients. The representations from each modality are projected into a 128-dimensional space before fusion, followed by a multilayer perceptron classifier with hidden-layer dimensions of 1024, 1024, 1024, and 512.

***Fine-Tuning Protocol.*** Each integrative model is optimized end-to-end, with both the RNA language model and MoLFormer fine-tuned rather than frozen. Training uses the Adam optimizer ($\beta_1 = 0.9$, $\beta_2 = 0.999$) with a weight decay of $10^{-6}$ as the $L_2$ penalty, a learning rate of $10^{-4}$ for every parameter group including the RNA language model, and a gradual warm-up schedule with a multiplier of 8 spanning the training budget. The objective is binary cross-entropy. Models are trained with a batch size of 32 for up to 100 epochs, with early stopping on validation AUROC and a patience of 20 epochs.

Each encoder is trained with three random seeds (1, 2 and 42) across the five cross-validation folds, yielding 15 trained models per encoder. Seven encoders are compared – LIMBER-*nano*, LIMBER-*macro*, RiNALMo-giga, AIDO.RNA-650M, RNA-FM, structRFM and RNASwan-seq – for 105 trained models in total. For RNASwan-seq, we do not use the published SMRTnet weights. Instead, the model is retrained from scratch under the same protocol, ensuring that its structural encoders are randomly initialized and trained under identical conditions to those used for all other models in the comparison.

***Ensemble Scoring on the Benchmark Dataset.*** For performance evaluation on the test folds, the 31-nucleotide binding-site fragments and their corresponding experimentally determined secondary structures are used directly as model inputs. The SMRTnet benchmark dataset does not provide such binding-site localization information; therefore, binding scores are obtained by systematically scanning each RNA sequence, as detailed in Algorithm2. Each RNA is scanned with 31-nucleotide windows at a stride of one nucleotide; every window, together with its experimentally resolved secondary structure, is scored by all five cross-validation models, and the median of the five predictions is taken as the window-level score. For RNAs of 31 to 40 nucleotides, the final score is the maximum over all windows. For RNAs longer than 40 nucleotides, candidate binding regions are identified as four consecutive windows that all score above 0.5, and the final score is the maximum among those four windows; if no such region exists, the minimum score across all windows is assigned, indicating a non-binding interaction. This procedure requires no prior knowledge of the binding site, at the cost of a local window design that may under-represent long-range interactions.

***Evaluation.*** On the cross-validation test folds, where the binding-site fragment and its secondary structure are supplied as inputs, performance is quantified by AUROC for each of the 15 trained models per encoder, and reported as the mean and standard deviation over those 15 values. On the benchmark dataset, scored by the ensemble procedure above, performance is quantified by accuracy, precision, recall, AUROC and AUPRC, each averaged over the three independent training seeds.

**Algorithm 2** Ensemble binding score for an RNA of unknown binding site

**Input:** RNA sequence $x$ of length $L \geq 31$ with secondary structure $s$; ligand SMILES $m$ and the molecular graph $g$ derived from it; the five cross-validation models $\{f_1, \ldots, f_5\}$; window length $w = 31$; run length $c = 4$; length cut-off $\ell = 40$

**for** $i = 1$ **to** $L - w + 1$ **do** ▷ all windows at stride 1
  $p_i \leftarrow \text{median}\big(f_1(x_{i:i+w-1}, s_{i:i+w-1}, m, g), \ldots, f_5(x_{i:i+w-1}, s_{i:i+w-1}, m, g)\big)$
**end for**
**if** $L \leq \ell$ **then**
  **return** $\max_i p_i$
**end if**
$R \leftarrow \{\, i \leq L - w - c + 2 \;:\; p_j > 0.5 \;\; \forall\, j \in [i,\, i + c - 1] \,\}$ ▷ starts of runs of $c$ consecutive windows above threshold
**if** $R \neq \emptyset$ **then**
  **return** $\max_{i \in R} \; \max_{j \in [i,\, i+c-1]} p_j$
**else**
  **return** $\min_i p_i$ ▷ no candidate region: scored as non-binding
**end if**

* These authors contributed equally to this work.

[1] Ashish Vaswani, Noam Shazeer, Niki Parmar, Jakob Uszkoreit, Llion Jones, Aidan N. Gomez, Lukasz Kaiser, and Illia Polosukhin. Attention Is All You Need, August 2023.

[2] Albert Gu, Karan Goel, and Christopher Ré. Efficiently Modeling Long Sequences with Structured State Spaces, August 2022.

[3] Albert Gu and Tri Dao. Mamba: Linear-Time Sequence Modeling with Selective State Spaces, May 2024.

[4] The RNAcentral Consortium. RNAcentral in 2026: Genes and literature integration. *Nucleic Acids Research*, 54(D1):D303–D313, January 2026.

[5] Padideh Danaee, Mason Rouches, Michelle Wiley, Dezhong Deng, Liang Huang, and David Hendrix. bpRNA: Large-scale automated annotation and analysis of RNA secondary structure. *Nucleic Acids Research*, 46(11):5381–5394, June 2018.

[6] Jaswinder Singh, Jack Hanson, Kuldip Paliwal, and Yaoqi Zhou. Rna secondary structure prediction using an ensemble of two-dimensional deep neural networks and transfer learning. *Nature Communications*, 10(1):5407, 2019.

[7] Elena Rivas, Raymond Lang, and Sean R. Eddy. A range of complex probabilistic models for RNA secondary structure prediction that includes the nearest-neighbor model and more. *RNA*, 18(2):193–212, January 2012.

[8] Kengo Sato, Manato Akiyama, and Yasubumi Sakakibara. Rna secondary structure prediction using deep learning with thermodynamic integration. *Nature Communications*, 12(1):941, 2021.

[9] Ioanna Kalvari, Eric P Nawrocki, Nancy Ontiveros-Palacios, Joanna Argasinska, Kevin Lamkiewicz, Manja Marz, Sam Griffiths-Jones, Claire Toffano-Nioche, Daniel Gautheret, Zasha Weinberg, Elena Rivas, Sean R Eddy, Robert D Finn, Alex Bateman, and Anton I Petrov. Rfam 14: Expanded coverage of metagenomic, viral and microRNA families. *Nucleic Acids Research*, 49(D1):D192–D200, January 2021.

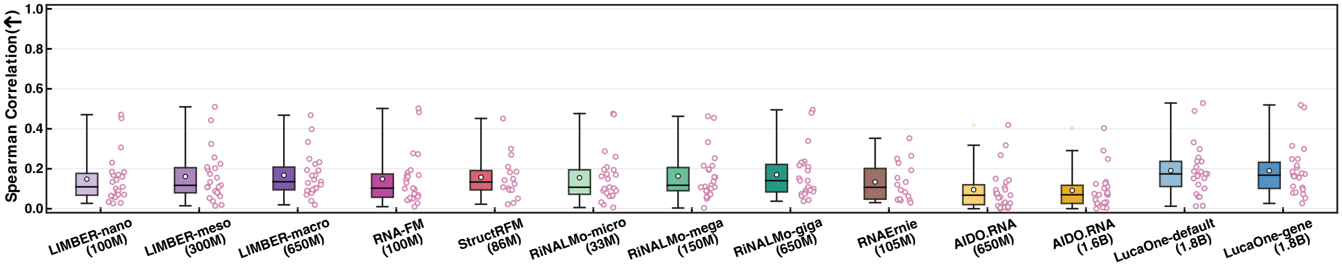


**Figure S1: Zero-shot fitness prediction of coding mRNAs by different RNA language models.** The absolute Spearman correlation between model-predicted relative log-likelihood scores and experimentally measured fitness or phenotypic effects is evaluated across 23 coding-mRNA deep mutational scanning assays. RNAGym sequences longer than 1,022 nucleotides are excluded. Each assay is represented by a point, while box plots summarize the overall distribution; circles and horizontal lines denote the mean and median, respectively.

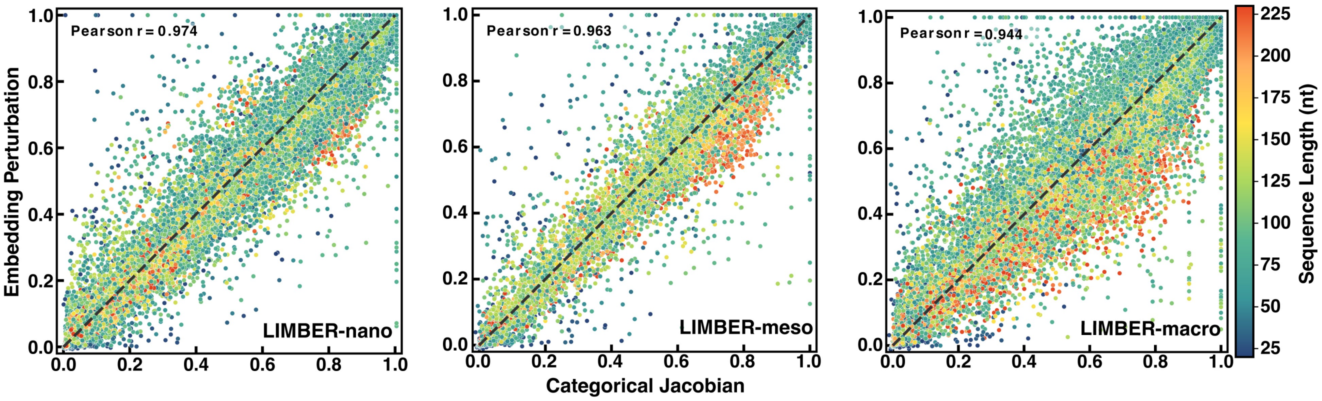


**Figure S2: Comparison of categorical Jacobian and embedding perturbation for zero-shot RNA secondary-structure prediction across LIMBER models with different sizes.** Secondary structures for sequences in the bpRNA-1m(90) dataset are inferred independently using the categorical Jacobian method (x-axis) and the embedding perturbation method (y-axis) with LIMBER-nano (left), LIMBER-meso (middle), and LIMBER-macro (right). Prediction accuracy for each sequence is quantified by normalized AUPR and plotted according to the corresponding values obtained from the two methods. Each point represents an individual RNA sequence and is colored by sequence length. The Pearson correlation between the two prediction approaches is calculated for each model size and reported in the corresponding panel.

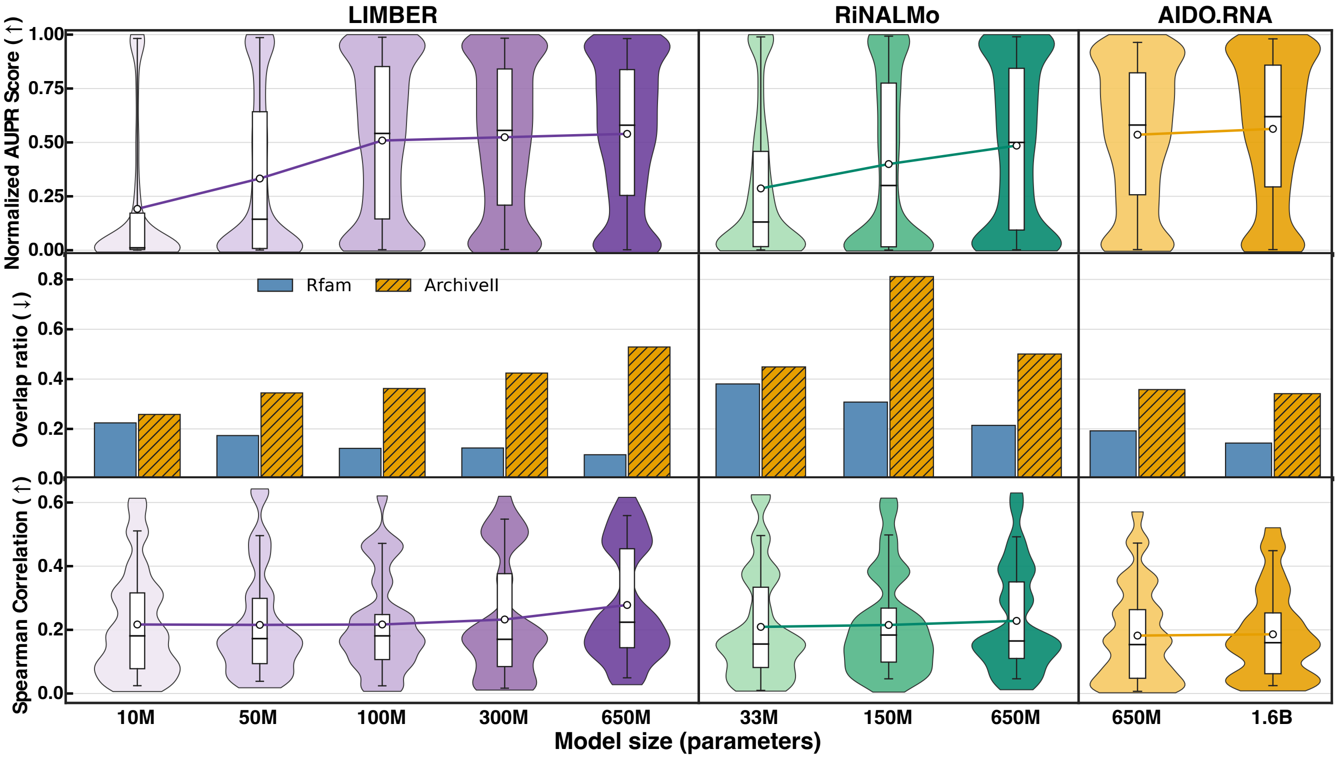


**Figure S3: Scaling of RNA language-model performance with model size across secondary-structure prediction, family classification, and fitness prediction.** Three RNA language-model families, i.e., LIMBER (10M, 50M, 100M, 300M and 650M), RiNALMo (33M, 150M and 650M), and AIDO.RNA (650M and 1.6B), spanning multiple model sizes are evaluated using zero-shot or unsupervised benchmarks. Top row: zero-shot secondary-structure prediction on the filtered bpRNA-1m(90) dataset, quantified by normalized AUPR from categorical-Jacobian-derived contact maps. Middle row: overlap ratios between homologous and non-homologous sequence representations in the final-layer embedding space for the Rfam and ArchiveII datasets. Bottom row: distributions of the absolute Spearman correlations between model-predicted log-likelihood scores and experimentally measured fitness or phenotypic effects across 33 deep mutational scanning assays. For all box plots, circles and horizontal lines denote the mean and median, respectively.

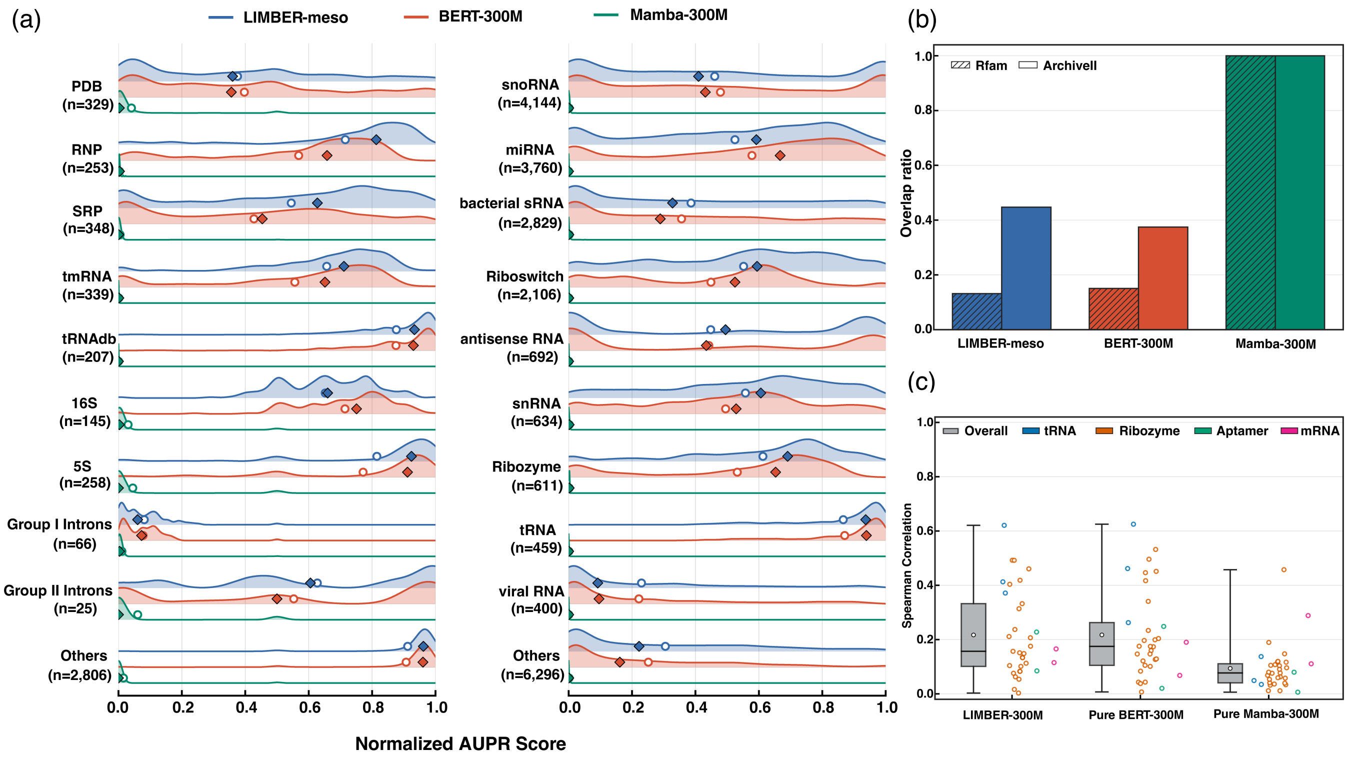


**Figure S4: Comparison of hybrid LIMBER with pure BERT and Mamba architectures across zero-shot benchmarks.** All models are matched in size ($\sim$ 300M parameters) and pretrained on the same dataset for six epochs (half the original training schedule). (a) Task-agnostic RNA secondary-structure recovery using categorical Jacobian analysis, evaluated by normalized AUPR. Mean and median scores are indicated by circle and diamond markers, respectively. (b) Discrimination between homologous and non-homologous RNA sequence pairs at the family (Rfam) and type (ArchiveII) levels, quantified by the overlap ratio of pairwise similarity distributions. (c) Zero-shot fitness prediction across 33 deep mutational scanning assays, evaluated by the absolute Spearman correlation between model-derived sequence likelihood scores and experimentally measured mutational fitness.

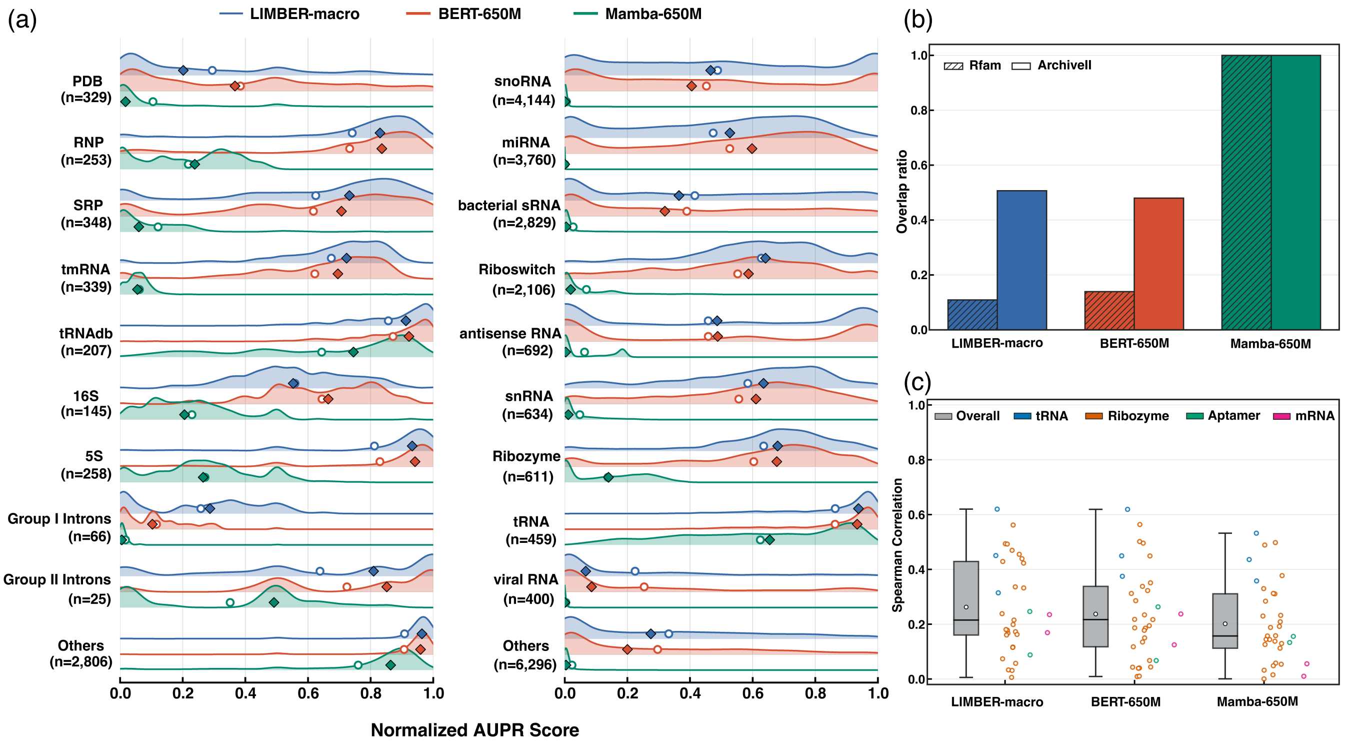


**Figure S5: Comparison of hybrid LIMBER with pure BERT and Mamba architectures across zero-shot benchmarks.** All models are matched in size ($\sim$ 650M parameters) and pretrained on the same dataset for six epochs (half the original training schedule). (a) Task-agnostic RNA secondary-structure recovery using categorical Jacobian analysis, evaluated by normalized AUPR. Mean and median scores are indicated by circle and diamond scattors, respectively. (b) Discrimination between homologous and non-homologous RNA sequence pairs at the family (Rfam) and type (ArchiveII) levels, quantified by the overlap ratio of pairwise similarity distributions. (c) Zero-shot fitness prediction across 33 deep mutational scanning assays, evaluated by the absolute Spearman correlation between model-derived sequence likelihood scores and experimentally measured mutational fitness.

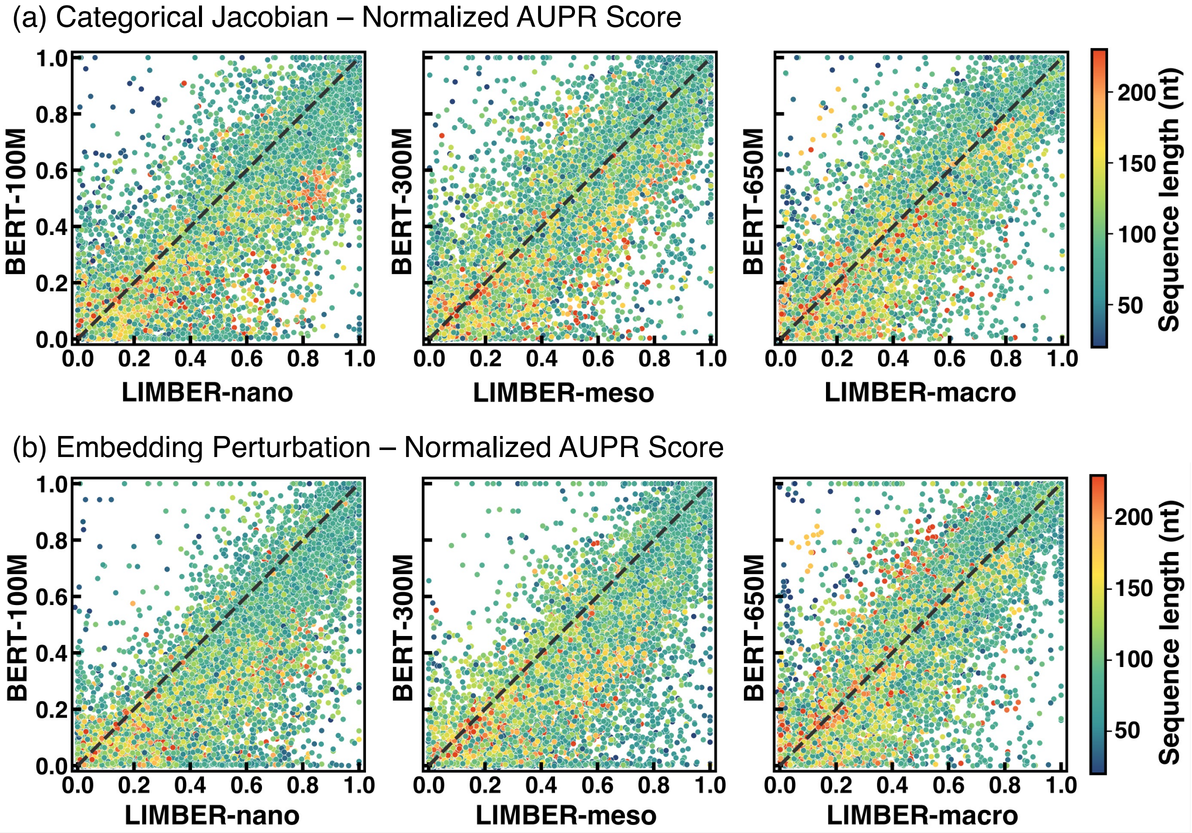


**Figure S6: Comparison of zero-shot secondary-structure recovery between hybrid LIMBER and pure BERT architectures across three model scales.** Two unsupervised approaches, categorical Jacobian and embedding perturbation, are used to infer nucleotide contact maps from model-output logits and final-layer embeddings, respectively, for more than 25,000 RNA sequences. Predicted contact maps are evaluated against reference secondary structures using normalized AUPR. Each point represents an individual RNA sequence, with the x- and y-axes indicating recovery performance for size-matched LIMBER and BERT models, respectively, and color denoting sequence length.

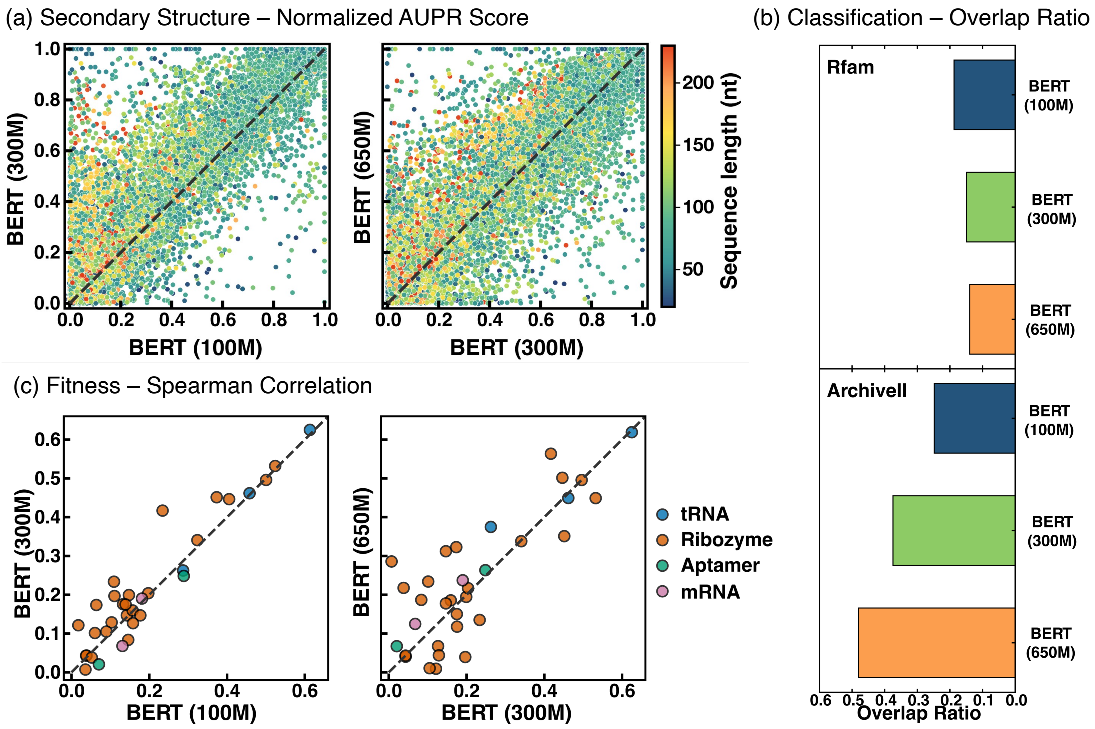


**Figure S7: Emergence of RNA structural and functional representations with BERT model scaling.** Zero-shot evaluations are compared across BERT models with 100M, 300M and 650M parameters. (a) Pairwise comparison of secondary-structure recovery across model scales using categorical Jacobian analysis, evaluated by normalized AUPR. Each point represents an RNA sequence, with the $x$- and $y$-axes indicating scores from smaller and larger models, respectively, and color denoting sequence length. (b) Zero-shot discrimination of homologous and non-homologous sequence pairs at the RNA-family (Rfam) and RNA-type (ArchiveII) levels, quantified by the overlap ratio of pairwise similarity distributions. (c) Pairwise comparison of zero-shot fitness prediction across model scales for 33 deep mutational scanning assays. Each point represents an assay, with the $x$- and $y$-axes indicating the absolute Spearman correlations between model-derived sequence likelihood scores and experimentally measured fitness for smaller and larger models.

(a)

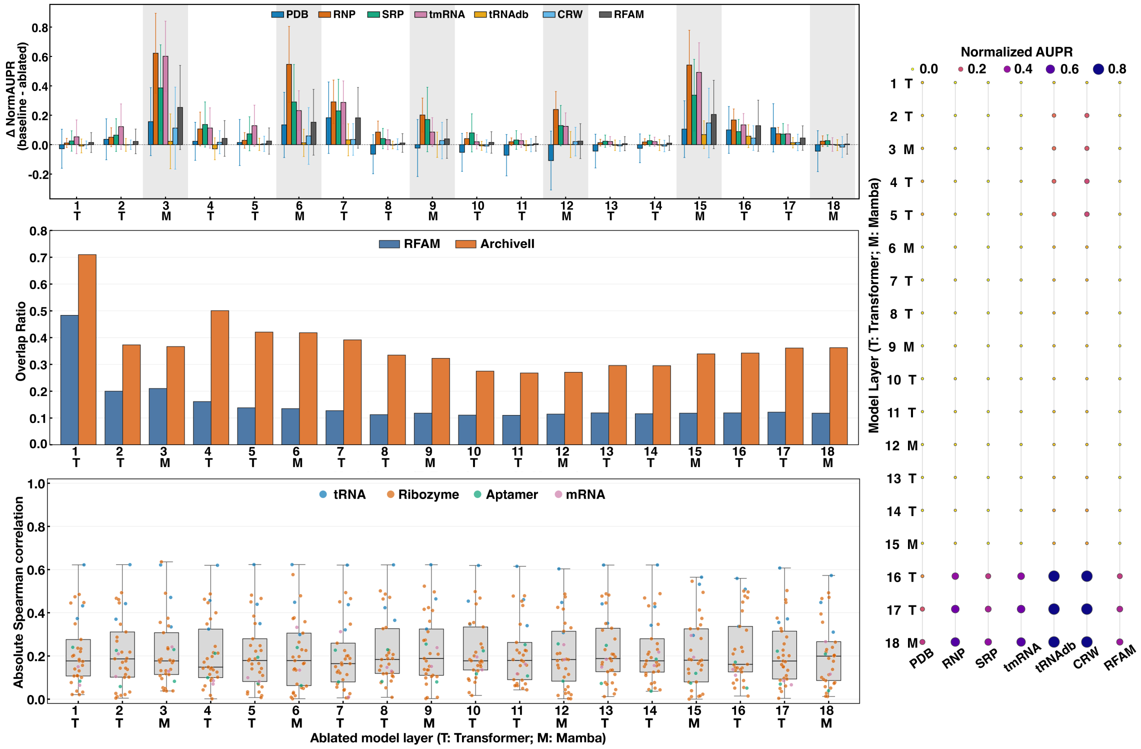


(b)

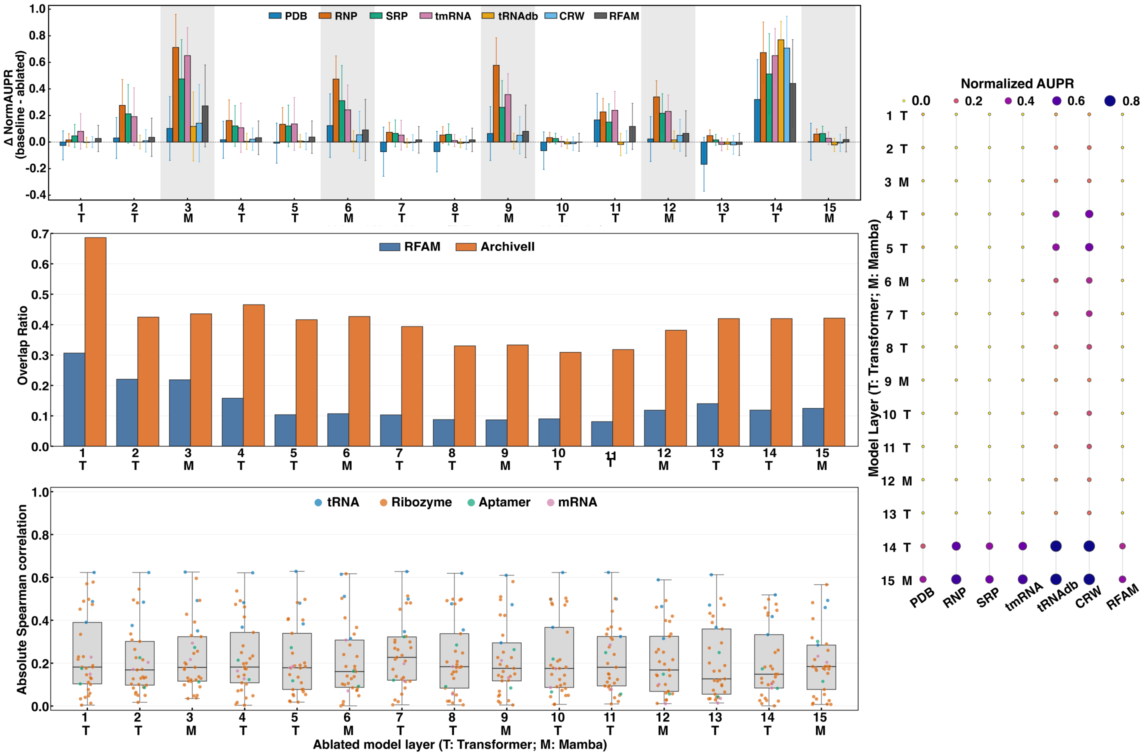


**Figure S8: Layer-wise Interpretation of LIMBER Internal Representations for RNA Structure Recovery, Classification, and Fitness Prediction.** Analyses are performed using (a) LIMBER-*nano* and (b) LIMBER-*meso*, both trained for 12 epochs, same as Fig. 4 of the main text.
